# A novel mouse model of *Campylobacter jejuni* enteropathy and diarrhea

**DOI:** 10.1101/283572

**Authors:** N Giallourou, GL Medlock, DT Bolick, PHQS Medeiros, SE Ledwaba, GL Kolling, K Tung, P Guerry, JR Swann, RL Guerrant

## Abstract

*Campylobacter* infections are among the leading bacterial causes of diarrhea and of ‘environmental enteropathy’ (EE) and growth failure worldwide. However, the lack of an inexpensive small animal model of enteric disease with *Campylobacter* has been a major limitation for understanding its pathogenesis, interventions or vaccine development. We describe a robust standard mouse model that can exhibit reproducible bloody diarrhea or growth failure, depending on the zinc or protein deficient diet and on antibiotic alteration of normal microbiota prior to infection. Zinc deficiency and the use of antibiotics create a niche for *Campylobacter* infection to establish by narrowing the metabolic flexibility of these mice for pathogen clearance and by promoting intestinal and systemic inflammation. Several biomarkers and intestinal pathology in this model also mimic those seen in human disease. This model provides a novel tool to testing specific hypotheses regarding disease pathogenesis as well as vaccine development that is currently in progress.

**Author Summary:** *Campylobacter jejuni* has been identified as one of the leading causes of enteropathy and diarrhea. In developing countries, these repeated enteric infections often result in growth deficits and cognitive impairment. There is a lack of small animal models of *Campylobacter* infection. This is a major hurdle in understanding the pathogenesis of *Campylobacter* infection in order to lead to therapeutic treatments and vaccines. We have developed a highly reproducible mouse model of *Campylobacter* infection that has clinical outcomes that match those of malnourished children. We hope that these insights into *Campylobacter* susceptibility will lead to the development of treatments against this major cause of diarrheal illness.

## Introduction

*Campylobacter jejuni* is a Gram-negative helical or rod shaped bacteria that is a common cause of gastroenteritis. *C. jejuni* is one of the leading recognized bacterial causes of food-borne illnesses in the U.S. and worldwide [1, 2].

In the Global Enteric Multicenter Study (GEMS) to identify the aetiology and population-based burden of pediatric diarrheal disease in sub-Saharan Africa and south Asia, *C. jejuni* was a leading bacterial cause of moderate to severe diarrhea burden at several South Asian sites (specifically in Bangladesh, Pakistan and India) [3]. In addition, *Campylobacter* spp. were the leading bacterial (and the leading pathogen overall in the second year of life) pathogen associated with diarrhea across all 8 sites in Asia, Africa and Latin America in the MAL-ED study [4]. Furthermore, “asymptomatic’ *Campylobacter* infections (i.e. without overt diarrhea; specimens collected at monthly surveillance) were also the leading pathogen associated with significant growth failure and with increased intestinal permeability and intestinal as well as systemic inflammatory biomarkers [5]. Finally, *Campylobacter* infections were significantly lower with exclusive breastfeeding, treatment of drinking water or by recent macrolide antibiotic use [5].

Classically, *Campylobacter jejuni* has been long recognized as a cause, much like *Salmonella* or *Shigella*, of febrile, often bloody and inflammatory diarrhea [6-8]. Diarrhea may also be accompanied by cramps and nausea, as well as headache and muscle pain. The incubation period is approximately 2-5 days after oral ingestion of contaminated water or food. *C. jejuni* diarrhea is typically self-limited to 7-10 days and relapses are uncommon. Unless the cause is recognized early, antibiotics are typically not prescribed although it is usually susceptible to macrolide antibiotics, including erythromycin and azithromycin that have been shown to reduce shedding of organism in stool and, if started early, the duration of symptoms [9].

While considered to be a microaerophilic organism requiring reduced levels of oxygen, *C. jejuni* can use oxygen in specific environments such as the chicken [10-14] cecum[15], in which it colonizes in a fucose-dependent manner and constitutes a major reservoir for human infections [16]. It is relatively fragile, and sensitive to environmental and oxidative stresses, as well as common disinfectants [17]. The organism requires 3 to 5% oxygen and increased (2 to 10%) carbon dioxide for optimal growth conditions. Prior to 1972, when methods were developed for its isolation from feces, it was believed to be primarily an animal pathogen causing abortion and enteritis in sheep and cattle [18]. This bacterium is now recognized as an important enteric pathogen [4, 19]. *Campylobacter* continues to be a leading recognized bacterial cause of diarrhea in the United States [20, 21].

While piglet, ferret, chick and even moth larvae have been studied as models, [22-26]; small animal models, such as mice, of *C. jejuni* have required genetic manipulations or have failed to exhibit enteric disease, limiting their benefit in translation to human disease [22, 27-29]. Stahl et al. has recently published several studies on *C. jejuni* infection in mice deficient in Single IgG Interleukin-1 related receptor (SIGIRR), a repressor of MyD88 [10-14]. An earlier model by McCardell et al. utilized iron-loaded BALB/C mice to elicit diarrhea within 4 hours after intraperitoneal injection of 10^9^ *C. jejuni* [30-32]. In this current study, we demonstrate a simple, affordable, and highly reproducible weaned mouse model of either enteropathy with growth failure or of overt bloody diarrhea, the two main recognized clinical manifestations of human disease with *Campylobacter* infections. Moreover, we demonstrate that, in addition to antibiotic disruption of resident microbiota, whether the mouse is fed a standard, zinc deficient, or protein deficient diet, has direct impact on the amount of colonization, shedding of organism in stool, inflammatory biomarkers, and intestinal architecture, as well as clinical outcomes that mimic human disease, including growth failure or overt, often bloody and inflammatory diarrhea.

## Results

### Antibiotics are required for Campylobacter jejuni colonization

Non-antibiotic treated C57BL/6 mice fed a standard house chow (HC) diet grew slightly better than antibiotic treated mice and showed similar weight loss following *Campylobacter* infection (Fig 1A). Both infected groups recovered weight quickly, 3 days post infection. However, there was a striking increase in *Campylobacter* shed in stool in antibiotic pretreated animals (Fig 1B). We were able to re-isolate and confirm the viable *C. jejuni* strain from feces of infected animals to confirm qPCR results (Supplemental Fig 1). Diarrhea was observed only in antibiotic pretreated infected animals. Additionally, the fecal biomarker myeloperoxidase (MPO) was significantly elevated only in the antibiotic-pretreated *Campylobacter* infected group (Fig 1C).

**Fig 1.**
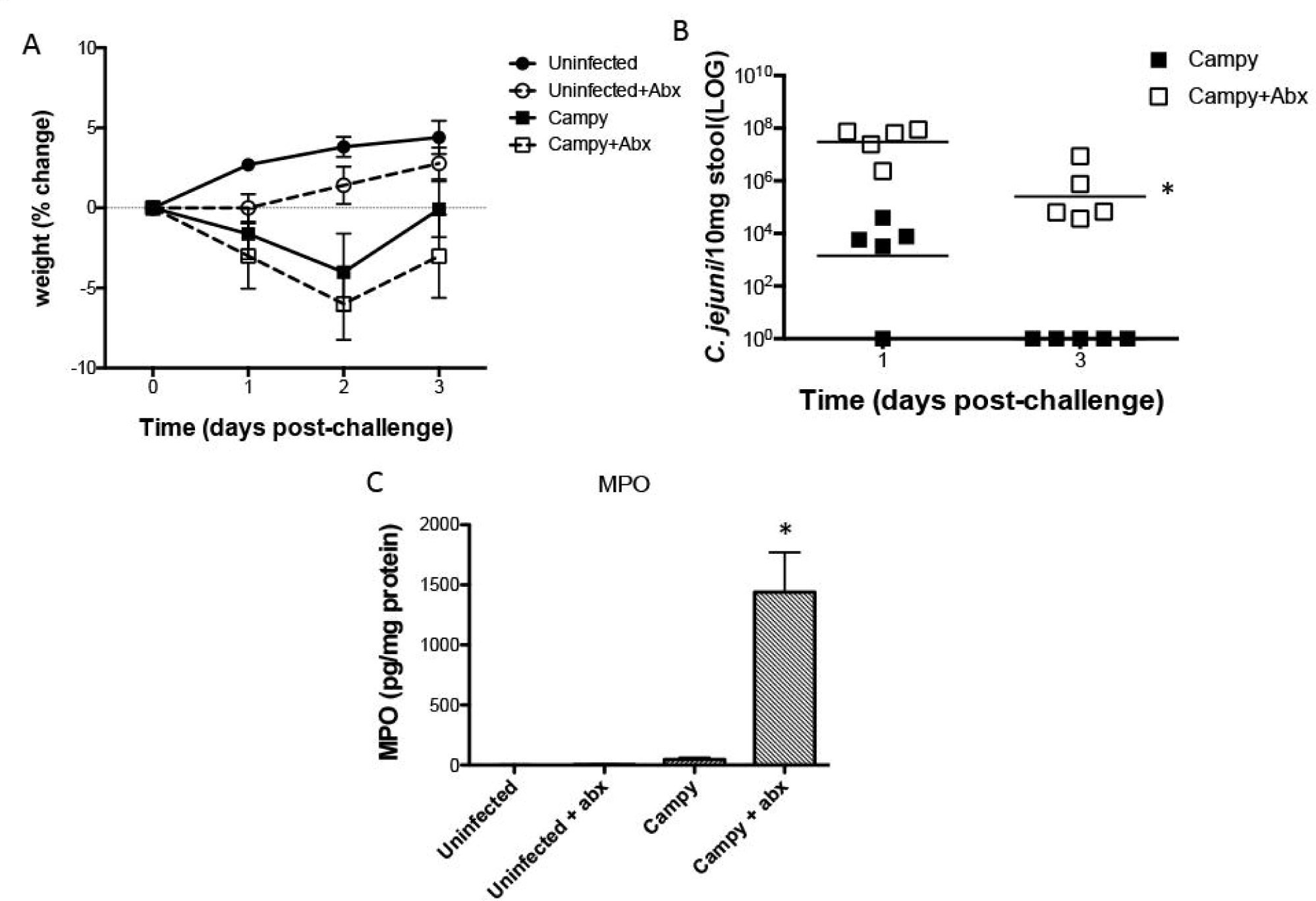
Antibiotics are required for *C. jejuni* colonization. Panel A. Weights following *C. jejuni* infection in house chow (HC) fed C57Bl/6J male mice. Panel B. qPCR detection of *C. jejuni* (*cadF*) following infection in house chow (HC) fed C57Bl/6J male mice. *, Significantly greater than no abx; P<0.05. Panel C. Myeloperoxidase (MPO) levels in mouse stool following *C. jejuni* infection. *, Significantly greater than all other groups; P<0.05. (N=5/group)

### Feeding diets deficient in protein or zinc alter Campylobacter outcomes

We have recently demonstrated very different outcomes of host metabolites and response to enteric infections depending on the nutritional status of the host [33-35]. Having established the benefit of antibiotic pretreatment in this model, we then probed the effect of zinc or protein deficiency in *Campylobacter* infection in antibiotic-pretreated mice. Mice (*n* = 8 / group) were infected with 10^7^ *C. jejuni* on day 14 following feeding of specific diet and antibiotic disruption of resident microbiota as described in methods. As shown in Fig 2A, infected mice fed a protein source diet without zinc (dZD) diet had significantly greater weight loss than infected mice receiving the HC diet or a diet deficient in protein (dPD). While all HC or dZD fed mice developed diarrhea by day 3 post infection, all dZD fed mice had persistent bloody diarrhea for greater than 7 days post infection (Fig 2C). No diarrhea was seen with mice fed the dPD diet. Only mice that were fed dZD diet were colonized with *C. jejuni* for the duration of the experiment as measured by shedding of organism in stool (Fig 2B). Similar to HC-fed mice, non-antibiotic treated dPD or dZD fed mice infected with *C. jejuni* did not develop diarrhea and cleared the organism by day 3 post infection (data not shown).

**Fig 2.**
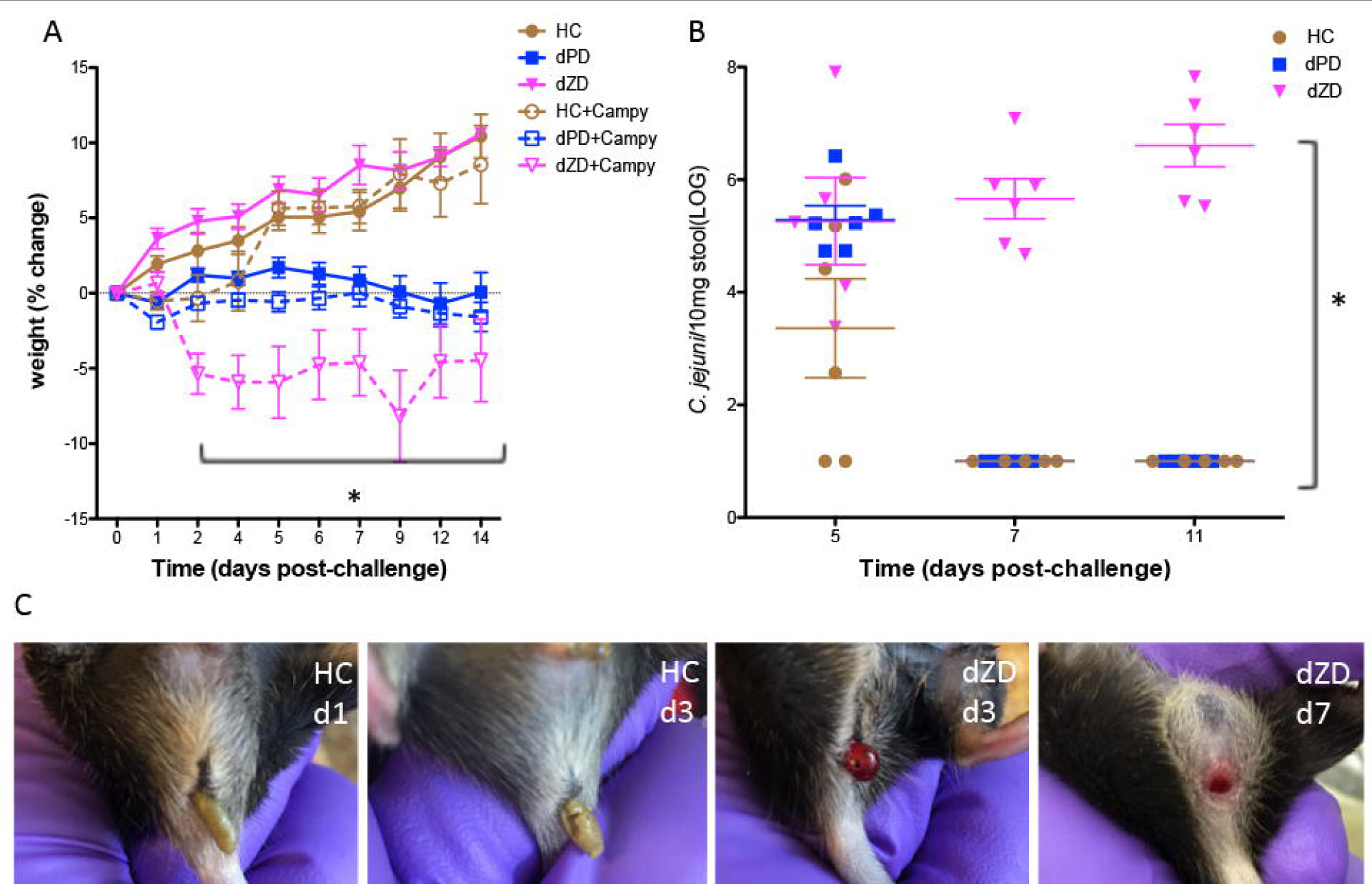
Prolonged weight loss following *Campylobacter* infection. Panel A. Mice fed either HC, or dPD had transient weight loss following infection, but dZD-fed infected mice had significant weight loss (* dZD infected vs uninfected days 2-14 post infection; P<0.001). While mice on dPD showed weight loss with no diarrhea, mice on HC had non-bloody soft stools on days 1-3 post infection, and mice on dZD had persistent bloody diarrhea on days 2-11 post infection. **Panel B.** Increasing Campylobacter detected in stool for the duration of the experiment (* dZD infected vs HC or dPD infected days 7&11 post infection; P<0.0001). **Panel C.** Images of stool following *Campylobacter* infection. Stool collected from HC (images from day 1 and 3 post infection) and dZD fed mice progressed from soft to bloody diarrhea by day 3 post infection (images from day 3 and 7 post infection). (N=8/group)

### Biomarkers of inflammation are increased following Campylobacter infection

MPO has been shown to increase with *Campylobacter* infections in humans across MAL-ED study sites [5]. In the MAL-ED studies, we demonstrated the correlation of MPO, lipocalin-2 (LCN-2), and calprotectin [36]. The fecal biomarkers of neutrophilic inflammation, MPO, and epithelial damage, LCN-2, were measured on day 2 and day 14 post *C. jejuni* infection. dZD-fed mice had significantly greater LCN-2 and MPO at both time points compared to control or protein deficient mice (Fig 3).

**Fig 3.**
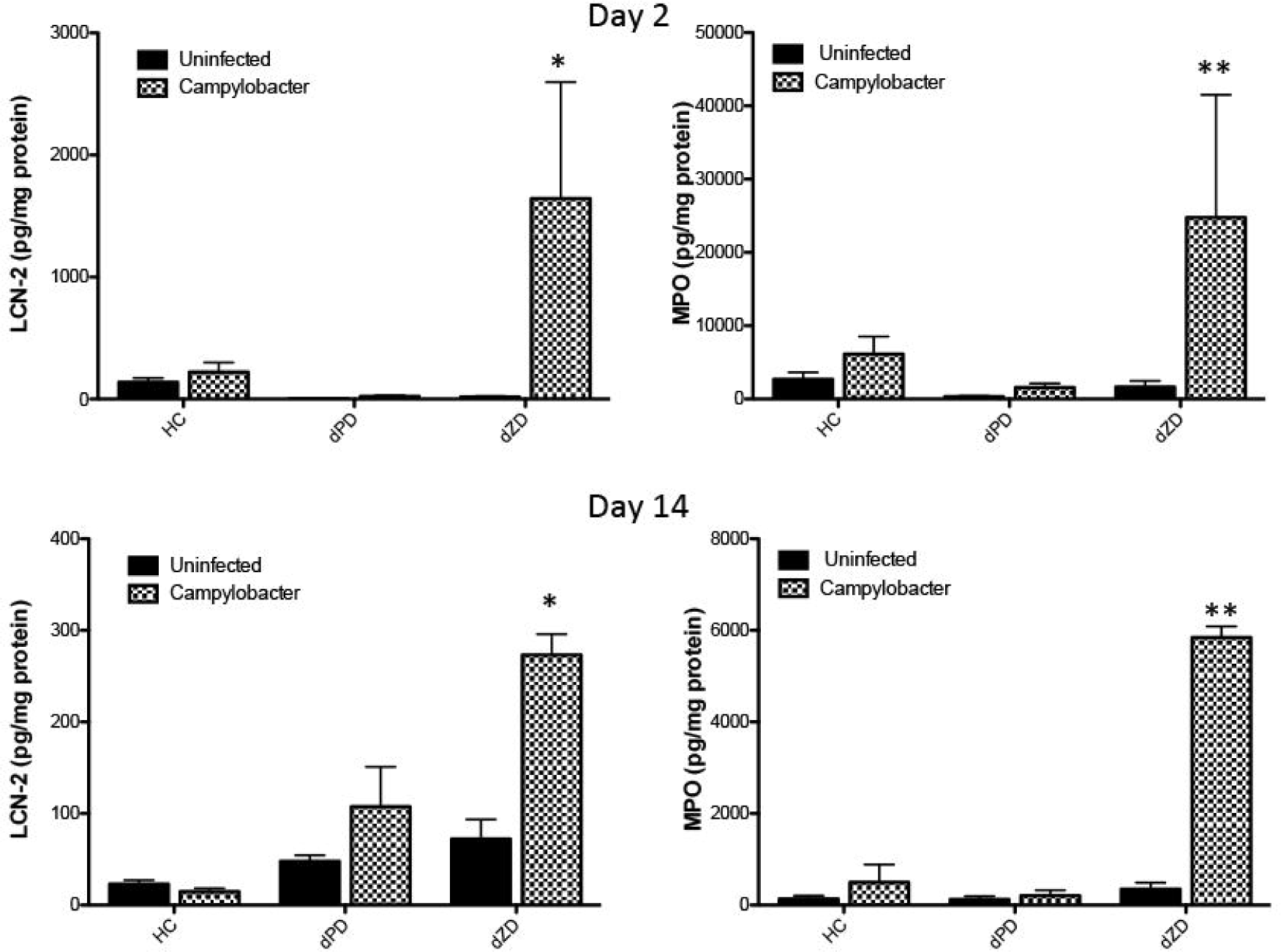
Inflammatory biomarkers following *Campylobacter* infection. The inflammatory biomarkers myeloperoxidase (MPO) and lipocalin-2 (LCN-2) were measured in cecal contents at day three post infection. LCN-2 was significantly elevated in dZD infected mice on both day 2 and 14 post infection, * dZD infected vs dZD uninfected; P<0.05. MPO was also significantly increased in dZD-fed infected mice on both day 2 and 14 post infection, ** dZD infected vs dZD uninfected; P<0.01.

### Intestinal morphology is altered by Campylobacter in zinc deficient mice

Finally, ileal histology of dZD-fed mice infected with *C. jejuni* showed increased mucus secretion and flattened villi (Fig 4A). As has been suggested in cecum by Stahl et al [29], we noted colonic submucosal edema and mucosal thinning, cell infiltration, crypt hyperplasia and striking goblet cell discharge (Fig 4B). These observations were confirmed by careful histopathologic assessment with a pathologist (KT). The predominant changes were seen in the colon, with increased lamina propria cellular infiltrate and striking lumenal mucus discharge (Supplemental Fig 2A), with only moderate mucus discharge seen in the ileum (Supplemental Fig 2B). In contrast, no any notable changes in histology were observed in HC or dPD-fed mice (data not shown).

**Fig 4.**
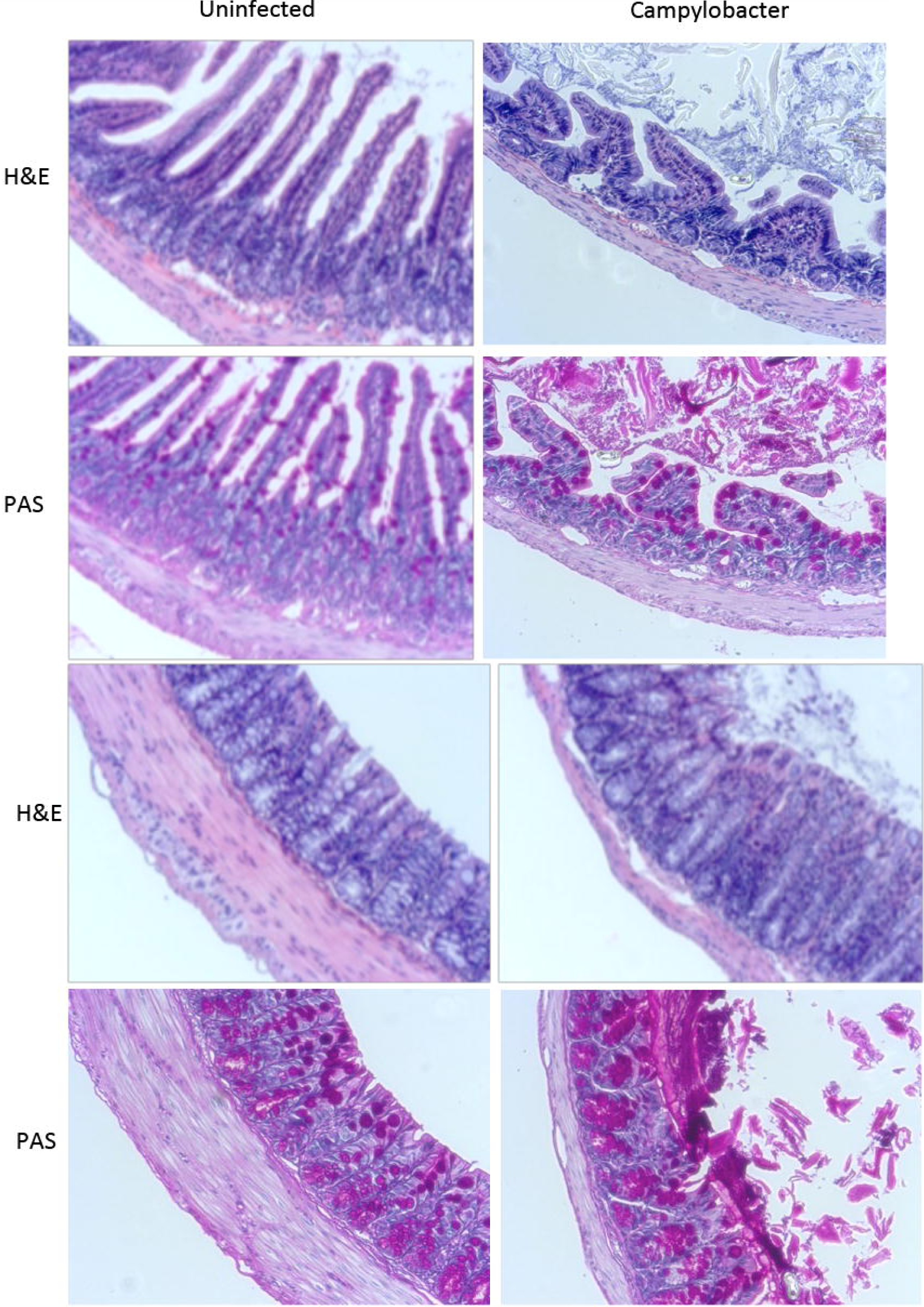
Intestinal morphometry. Intestinal sections from dZD fed mice stained for H&E or PAS. **Panel A:** ileum. **Panel B:** colon.

### Campylobacter capsule gene kpsM is required for colonization

Having established that zinc deficiency greatly enhanced *C. jejuni*, we then tested a non-encapsulated *kpsM* mutant of *C. jejuni* 81-176 [37]. All mice were fed the dZD diet and pretreated with antibiotics to disrupt resident microbiota as above. Mice infected with the *kpsM C. jejuni* grew significantly better than wildtype infected mice (Fig 5A). Additionally, shedding of *C. jejuni kpsM* was undetectable following day 1 suggesting an inability to colonize as the wildtype strain (Fig 5B). Diarrhea was observed only in wildtype, infected mice.

**Fig 5.**
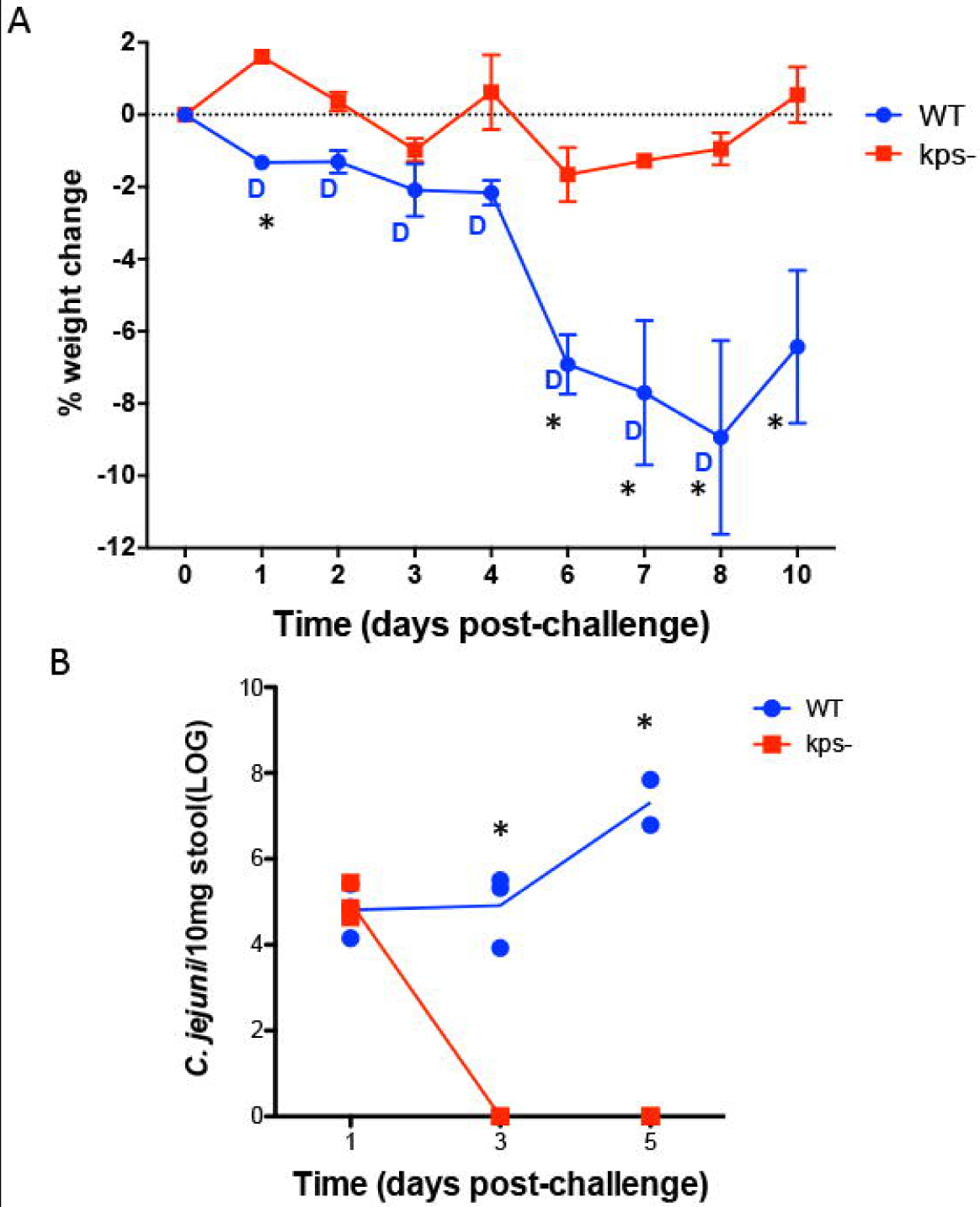
A non-encapsulated mutant of *C. jejuni* is defective in weight loss, diarrhea, and shedding of organism in stool. Panel A. Weights following infection with either wildtype or 81-176 *kpsM* in dZD C57Bl/6J male mice. * WT vs *kpsM*, P<0.05. Diarrhea noted by D. **Panel B.** qPCR detection of *Campylobacter jejuni* (*cadF*) following infection in dZD mice. * WT vs *kpsM*, P<0.05. (N=4/group)

### Metabolic perturbations induced by C.jejuni infection

Orthogonal projection to latent structures-discriminant analysis (OPLS-DA) models were constructed to assess the urinary metabolic perturbations induced by the *Campylobacter jejuni* challenge on days 5, 9, 11 and 14 post-infection. No significant differences in the urinary metabolic profiles of infected and uninfected mice fed the protein deficient diet were observed, consistent with the quick clearance of the pathogen and absence of diarrhea. Nourished mice differed from their infected counterparts on day 2 post-infection (Q^2^Y= 0.42, P = 0.04). Nourished mice were observed to excrete less taurine in the presence of the infection at this sampling point.

In agreement with results obtained from the growth, inflammatory and shedding measures, the most notable metabolic changes in response to infection were seen in dZD-fed mice. Significant metabolic changes in zinc deficient uninfected and infected mice are summarized in Fig 6 along with their correlation to class.

**Fig 6.**
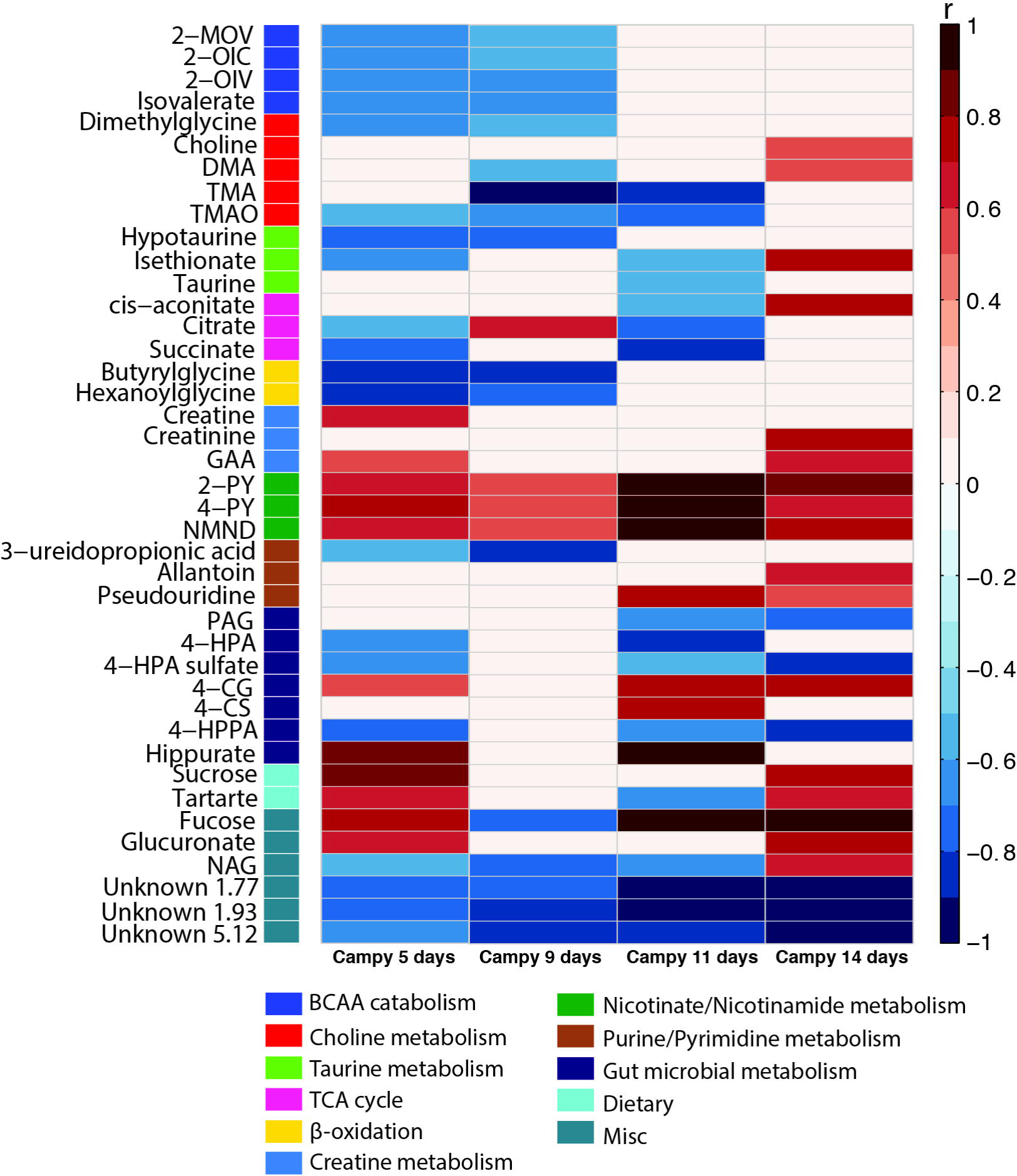
Metabolic perturbations induced by *Campylobacter* infection. Heatmap of the significant metabolic perturbations induced by C.jejuni infection identified by OPLS-DA models. Metabolic shifts are represented as correlation coefficients (r) of infected mice at days 4, 5 and 9 post infection versus diet and age-matched uninfected animals. Red and blue colors indicate increased or decreased excretion of metabolites following C.jejuni challenge respectively. Abbreviations: 2-OIV, 2-oxoisovalerate; 2-OIC, 2-oxoisocaproate; 2-MOV, 3-methyl-2-oxovalerate; 2-PY, N-methyl-2-pyridone-5-carboxamide; 4-CG; 4-cresol glucuronide; 4-CS, 4-cresyl sulfate; 4-HPA, 4-hydroxyphenylacetate; 4-HPPA, 4-Hydroxyphenylpyruvate; 4-PY, N-methyl-4-pyridone-3-carboxamide; DMA, dimethylamine; GAA guanidinoacetate; NAG, N-acetyl glutamine; NMND, N-methylnicotinamide; PAG, phenylacetylglycine; TMA, trimethylamine; TMAO, trimethylamine-N-oxide.

Following 5 (Q^2^Y= 0.58, P = 0.018) and 9 days (Q^2^Y= 0.73, P = 0.003) of *C. jejuni* infection mice excreted lower amounts of metabolites resulting from the catabolism of branched-chain amino acids [3-methyl-2oxovalerate (2-MOV), 2-oxoisocaproate (2-OIC), 2-oxoisovalerate (2-OIV), isovalerate], as well as glycine conjugates of fatty acid β -oxidation (butyrylglycine, hexanoylglycine). Differences in the excretion of these metabolites between uninfected and infected dZD mice are not present at day 11 (Q^2^Y= 0.82, P = 0.023) and 14 (Q^2^Y= 0.56, P = 0.028) post-infection. Following the *C. jejuni* challenge, the downstream metabolites arising from the systemic metabolism of tryptophan and kynurenine (*N*-methylnicotinamide (NMND), *N*-methyl-2-pyridone-5-carboxamide (2-PY), *N*-methyl-4-pyridone-3-carboxamide (4-PY)) were excreted in greater amounts at all sampling points. This is consistent with a sustained attempt to dampen the inflammatory response to infection through the induction of indoleamine 2,3-dioxygenase.

Additionally, shifts in the excretion of gut microbial-host co-metabolites were observed on days 5, 11 and 14 post-infection. Phenylacetylglutamine (PAG), a co-metabolite derived from phenylalanine, and 4-hydroxyphenylacetate (4-HPA) and 4-HPA sulfate, derivatives of tyrosine, were excreted in lower amounts in the *C. jejuni* infected mice. 4-HPA can be further processed by intestinal bacteria to 4-cresol, which once absorbed is either glucuronidated or sulfated in the liver to 4-cresyl glucuronide (4CG) and 4-cresyl sulfate (4CS), respectively prior to excretion. Both 4-CG and 4-CS were excreted in greater amounts in the infected dZD mice compared to their uninfected equivalents.

Consistent with histopathological findings, an increase in the excretion of diet-derived sucrose and tartrate was observed at days 5 and 14 post-infection indicative of impaired barrier function. Changes in the excretion of metabolites associated with oxidative stress and intestinal damage like allantoin, pseudouridine and fucose, were also noted in the infected animals. These findings are in agreement with the elevated biomarkers of intestinal inflammation and disturbed gut morphology in infected mice fed the dZD diet. Correlation analysis between biomarkers of inflammation and urinary metabolites identified significant positive correlations between LCN-2 and fucose and between MPO and hexanoylglycine and the two gut microbial derived metabolites 4-CS and hippurate in zinc deficient infected mice (Fig 7).

**Fig 7.**
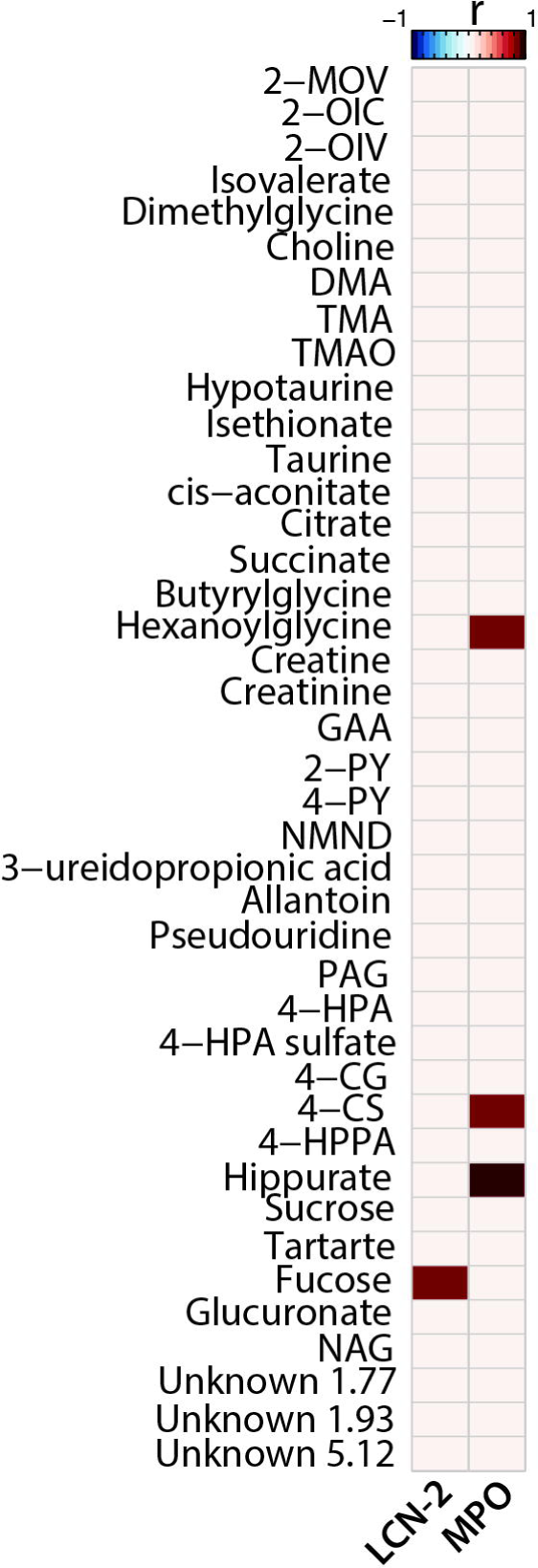
Correlations between urinary metabolites and inflammatory biomarkers. Spearman’s correlation heatmap between the urinary metabolites identified in the OPLS-DA models and the levels of inflammatory biomarkers LCN-2 and MPO on day 14 post infection. Only significant correlations following *P* value adjustment are shown (Benjamini-Hochberg for 5% false discovery rate).

### Perturbations to the microbiota induced by C.jejuni infection

16S rRNA gene sequencing was performed on fecal samples collected from infected and uninfected mice on all diets at 0, 1, 5, and 9 days post-infection. Richness (defined here as the number of unique amplicon strain variants (ASVs) detected in each sample) and evenness (the inverse of the Simpson index) of the microbiota was calculated for all samples (Fig 8A and 8B). Infected dPD-fed mice had higher richness at day 5 post-infection than at all other time points for both infected and uninfected dPD-fed mice. Similarly, infected dPD-fed mice at day 5 post-infection had higher evenness than any other time point for both infected and uninfected dPD-fed mice. Uninfected dPD mice had higher evenness at day 9 post-infection than all dPD samples other than those from infected mice at days 5 and 9 post-infection.

**Fig 8.**
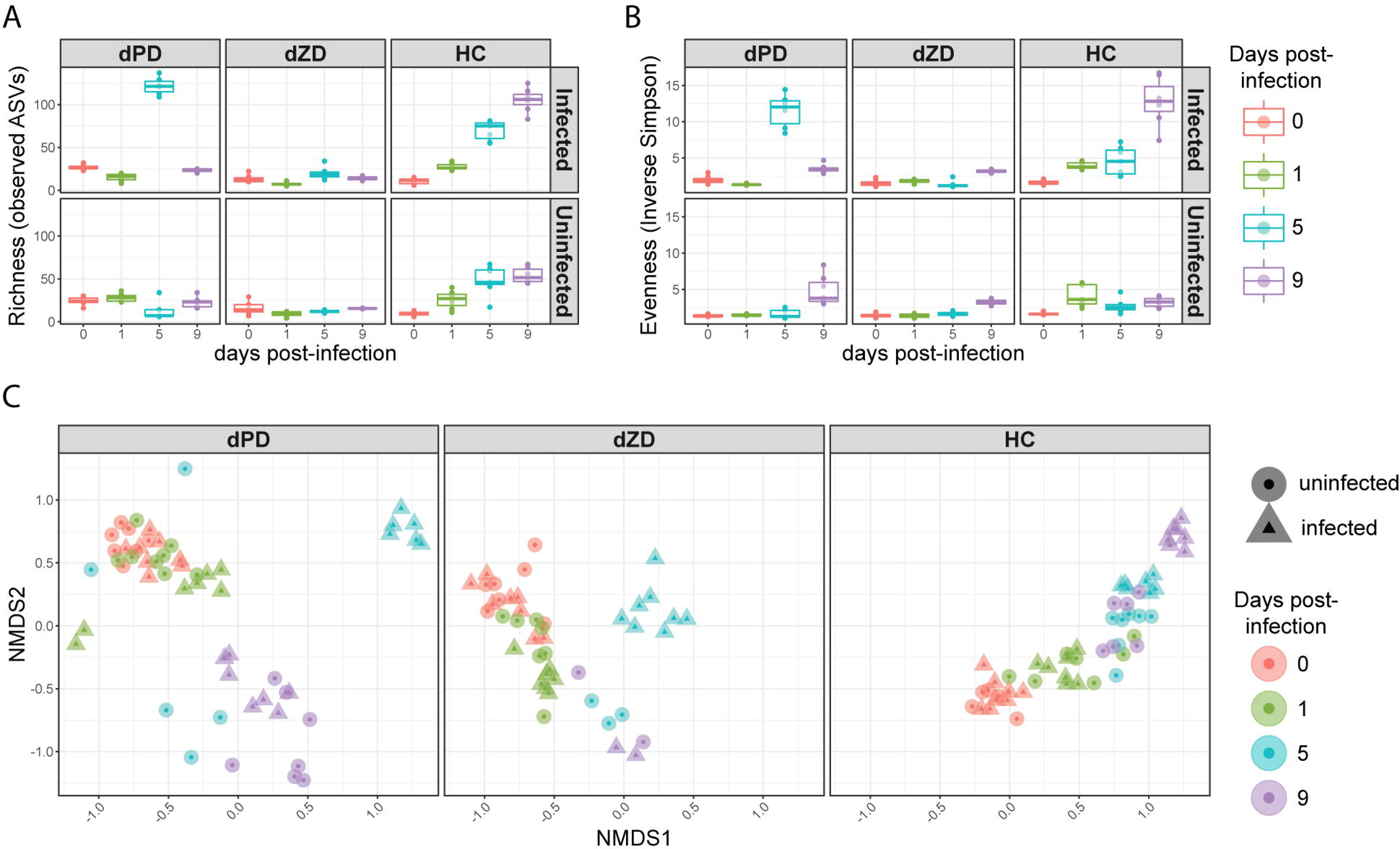
Summary of diversity and composition of the fecal microbiota of infected and uninfected mice on each diet. A) Richness (number of unique amplicon sequence variants (ASVs)) in each sample. B) Evenness (inverse of the Simpson Index) for each sample. C) Non-metric multidimensional scaling (NMDS) of the bray-curtis dissimilarity of each sample. NMDS was performed using all samples shown in the three panels, then samples were plotted separately by diet to improve visualization.

dZD-fed mice had no significant differences in richness or evenness over time within infection status or across infection status. In infected HC-fed mice, richness was lower at 0 days post infection than at all other time points for infected and uninfected HC-fed mice. The same was true for uninfected HC-fed mice, except that no significant difference was detected in the comparison with uninfected HC-fed mice at 1 day post-infection. No significant difference was detected in richness between uninfected and infected HC-fed mice at 0 days post-infection. In infected HC-fed mice, richness increased with each successive time-point. The same was true for uninfected HC-fed mice, except that richness plateaued at 5 days post-infection. Evenness was higher in infected HC-fed mice at 9 days post-infection than at all other time points for both infected and uninfected HC-fed mice. Evenness of infected HC-fed mice at day 0 was lower than at all other time points.

We calculated beta diversity to describe the global dissimilarity of all samples in the experiment (Fig 8C). Within each diet, uninfected and infected samples from the same time point tended to cluster together, except for dPD- and dZD-fed mice at 5 days post-infection and HC-fed mice at 9 days post-infection. For dPD-fed mice, this time point is also when the transient increase in richness and evenness occurs, and the samples are compositionally similar to the infected HC-fed mice at 9 days post-infection.

Given that mice were on antibiotics prior to infection with *C. jejuni*, the increases in richness and evenness over time in HC-fed mice are expected. Surprisingly, infected HC-fed mice had higher evenness and richness than uninfected mice at 9 days post infection. Antibiotic cessation may not increase evenness and richness in dPD- and dZD-fed mice given the incomplete diets they are receiving, from which dietary fiber is also absent. Substantial increases in either richness or evenness over time were not observed for either defined diet with or without infection. However, infected dPD-fed mice had a transient increase in evenness and richness at 5 days post infection, which is also the last time point for which shedding is detected from dPD-fed mice. Infected dPD-fed mice at this time point had higher abundance of amplicon sequence variants (ASVs) assigned to Bacteroidales S24-7 group, several from the order Clostridiales,,multiple *Enterobacteriaceae*, as well as several other ASVs (Fig 9). Many of these increases persisted to day 9 post-infection. Overall, infection led to a greater number of differentially abundant ASVs in dPD-fed mice at days 5 and 9 post-infection than any other diet or time point.

**Fig 9.**
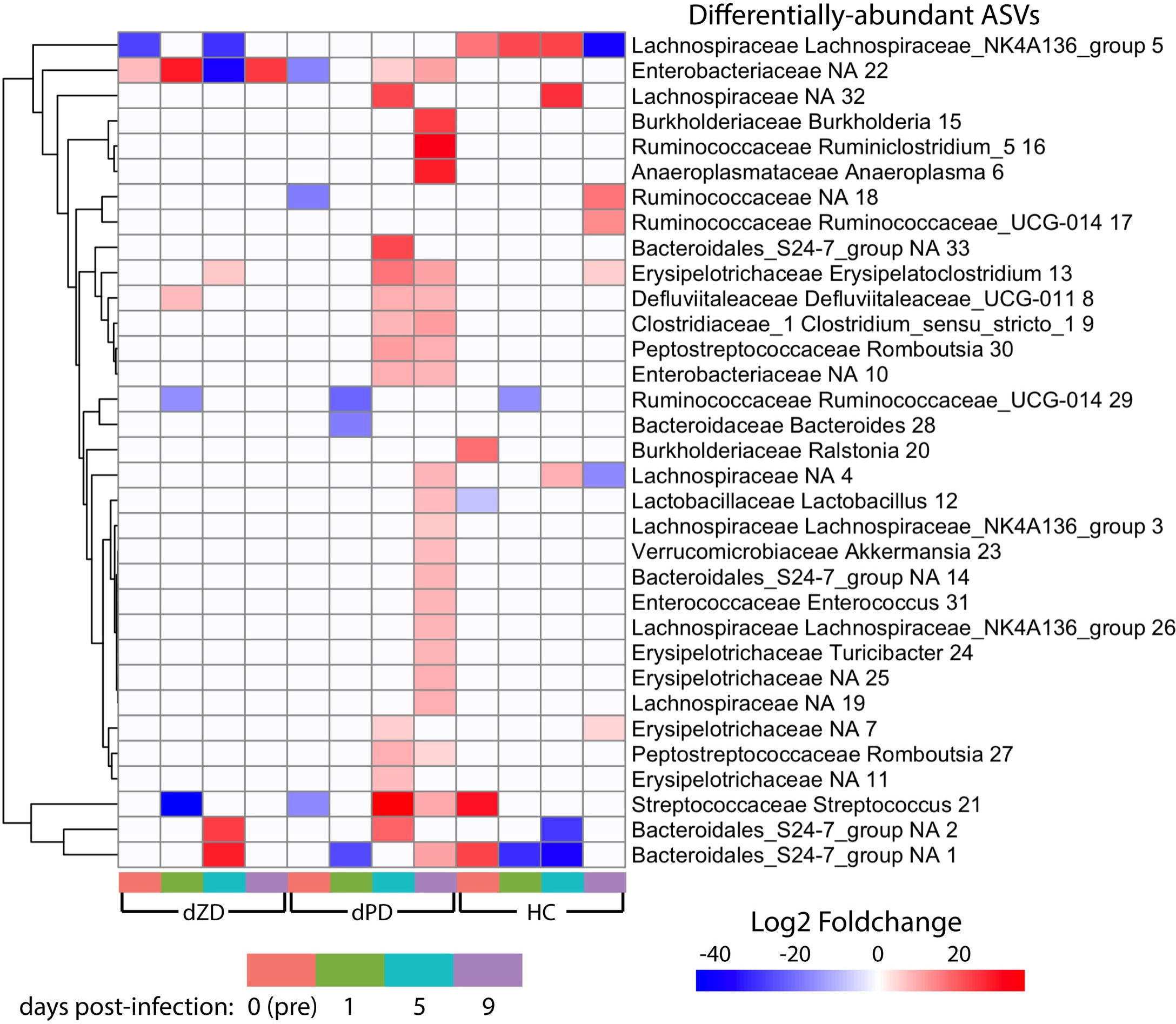
**Heatmap of differentially-abundant ASVs**. For each time point within each diet, microbiota were compared between infected and uninfected samples. Differentially-abundant ASVs from any comparison are shown in the heatmap, and the color of each cell indicates the log2 fold-change calculated with DESeq2. For each condition, only ASVs with a multiple testing-corrected p < 0.05 are shown (otherwise, cell is white). ASV labels include assigned family, genus and an ASV number used to differentiate multiple ASVs with identical genus-level assignment.

Infection had varied effects on the microbiota of dZD-fed mice over time. At day one post-infection, there was an increase of ASVs assigned to *Enterobacteriaceae*, *Defluviitaleaceae* UCG-011, and decreases in ASVs assigned to *Ruminococcaceae* UCG-014 and *Streptococcus* (Fig 9). This change is consistent with blooms of *Enterobacteriaceae* observered immediately after infection with other pathogens that may be induced by the host inflammatory response [38]. However, none of these changes persisted through day 5 post-infection, where the same *Enterobacteriaceae* ASV was substantially less abundant in the infected group rather than more abundant as at day 1 post-infection. Two ASVs assigned to the S24-7 group of the Bacteroidales order and an ASV assigned to the genus *Erysipelatoclostridium* were increased in infected mice, while *Lachnospiraceae* NK4A135 decreased. By day 9 post-infection, no differences were observed between infected and uninfected dZD-fed mice other than in increase in *Enterobacteriaceae* in infected mice. Given that dZD-fed mice had persistent bloody diarrhea, the relatively low number of differentially-abundant ASVs is may be due to their extremely limited diversity, with fewer than 25 unique ASVs in all but two samples at all time points.

HC-fed mice also had few ASVs that were consistently differentially-abundant over time. Two ASVs assigned to the S24-7 group of the Bacteroidales order were decreased in abundance at day 5 post-infection, one of which was also decreased at day 1 post-infection. At day 1 post-infection, one *Ruminococcaceae* UCG-014 ASV was also decreased in abundance, while *Lachnospiraceae* NK4A135 increased. At day 5 post-infection, the change in *Lachnospiraceae* NK4A135 persisted and two additional *Lachnospiraceae* ASVs had increased abundance. At day 9 post-infection, *Lachnospiraceae* NK4A135 and one additional *Lachnospiraceae* ASV were less abundant in infected mice than uninfected mice, and 2 Ruminococcaceae ASVs and 2 Erysipelotrichaceae ASVs were more abundant in infected mice.

## Discussion

Given the increasingly recognized importance of *Campylobacter* infections (as causes of severe bloody diarrhea or growth failure in children living in developing areas) and of food and milk-borne outbreaks in developed areas, not to mention the troubling sequelae like Guillain-Barré Syndrome (GBS), the need for improved understanding of its pathogenesis and effective interventions or vaccines is paramount. A major limitation of work in this area is the lack of an inexpensive animal model that mimics human disease. Here, we demonstrate that nutritional and antimicrobial manipulations can enable us to model either ‘asymptomatic’ enteropathy and growth failure or overt bloody diarrhea. This model opens tremendous opportunities to help enable vaccine development or other potential interventions.

The biomarkers that are common to both humans and our murine models also provide relevant markers to assess these interventions and mechanisms as well as the directly relevant clinical disease outcomes [5, 36]. Interestingly, a non-encapsulated mutant of 81-176 was severely attenuated in colonization and disease in animals fed the zinc deficient diet. This same mutant was attenuated in a ferret model of diarrheal disease and was slightly reduced in its ability to colonize BALB/c mice compared to the 81-176 parent [37, 39]. Although not shown in the current study, older mice (>10 weeks of age) were susceptible to *C. jejuni* infection; hence this model should enable vaccine studies. Whether we can model post-infectious sequelae such as GBS [40] or the roles of the microbiome in susceptibility to disease or sequelae remains to be seen.

There are several limitations on any non-human model of infectious diseases that must always be kept in mind. Coprophagic rodents clearly have quite different microbiota, diets and host determinants of disease. Nevertheless, recognizing these limitations, the extent to which we can model key major disease outcomes, including shared biomarker and even metabolomic pathways [35, 36, 41, 42] in an inexpensive, relative to existing models, defined murine model (without major genetic alterations or artificial pharmacologic immune manipulation) at least enables hypothesis generation and testing, as well as potential new vaccines that are currently in development [20].

*C.jejuni* infection in the zinc deficient mice elicited a robust inflammatory response evident by bloody diarrhea, intestinal and systemic inflammation parallel to growth faltering, highlighting the direct relevance of this model in clinical Campylobacteriosis research. This was accompanied by an activation of the kynurenine pathway as a result of systemic inflammation in response to infection. Indoleamine 2,3-dioxygenase (IDO) metabolizes tryptophan to kynurenine and it is increased in response to infection and inflammation resulting in a greater excretion of NMND and its downstream metabolites 2-PY and 4-PY. NMND is generated from nicotinamide adenine dinucleotide (NAD) by the action of the enzyme *N-*methyl transferase (NMNT), which is considered a master regulator of energy expenditure [43]. Increased excretion of NMND by zinc deficient infected mice reflects lower energy expenditure. The energy acquisition and utilization of the host is compromised by zinc deficiency [44] (Supplemental Fig 3) and further exacerbated following *C.jejuni* infection. Metabolic exhaustion suggests reductions in the levels of available NAD and here is reflected in the compromised activity of the energy generating TCA cycle (decreased citrate and succinate excretion) that happens concurrently to the loss of β-oxidation activity (hexanoylglycine and butyrylglycine), branched-chain amino acid catabolism (2-MOV, 2-OIC, 2-OIV) and muscle catabolism (increased creatine excretion on day 5 post infection). Zinc is a necessary cofactor for numerous enzymes involved in energy regulation [45], zinc deficiency therefore narrows the metabolic flexibility of the infected mice hindering pathogen clearance. We invite parallels to be drawn between the metabolic phenotype of the *C.jejuni* infected mice described here and that of our murine model of *Shigella flexneri* (manuscript currently under peer-review) also achieved with zinc deprivation and the use of antimicrobials. Zinc deficient *S.flexneri* infected mice displayed a similar metabolic profile characterized by energy exhaustion and catabolic stress combined with delayed pathogen clearance in these mice, compared to the HC-fed mice.

The presence of *C. jejuni* in the gut of zinc deficient mice perturbed metabolites derived from the gut microbiota. While PAG, 4-HPPA, 4-HPA and 4-HPA sulfate excretion was decreased with infection; the glucuronidated and sulfated forms of 4-cresol (microbial metabolites of 4-HPA) were increased. 4-cresol is a known bacteriostatic, negatively influencing the growth of gut bacteria [46] contributing to the lack of microbial diversity in the zinc deficient infected mice. Interestingly, increased urinary excretion of 4-CS was positively correlated with the fecal inflammatory biomarker, MPO. Increased excretion of 4-CS implies greater production and exposure of 4-cresol in the gut. This suggests that the release of 4-cresol in the intestine may contribute to the acute inflammatory response and subsequent damage to the intestinal epithelium. Although further work is necessary to establish causality. The lack of persistent changes in microbiota composition in dZD-fed mice make any signature difficult to resolve. Metagenomic studies are needed to identify the key functions disrupted by infection, as well as biomarkers of infection. The dynamics of differentially-abundant ASVs over the course of infection suggest that microbiota composition changes more rapidly as a result of infection than the host metabolome. In the other diets (dPD and HC), the ASVs that become differentially-abundant upon infection have little overlap with those from dZD-fed mice. This implies that the effect of infection on the microbiota is diet-dependent, which could be due to the difference in microbiota background induced by the diet, or that the intestinal environment changes in unique ways with each phenotype resulting from infection while being fed each diet. It seems likely that a combination of these factors (e.g. pathogen-microbiome, diet-pathogen, host-diet, host-microbiome, and microbiome-diet interactions) influence the virulence of the pathogen. It will be interesting to compare metabolic responses with *C. jejuni* infection in mammals with those in the avian reservoir(s), in which intriguing effects of microbial metabolites associated with reduced colonization after potential vaccine and probiotics has recently been reported [47].

Intriguingly, the glycoprotein marker of epithelial damage LCN-2 positively correlated with the excretion of fucose. Fucose is a sugar, found in high quantities in the mammalian gut either as a part of glycan structures or as a free monosaccharide and current research suggests it can have pleiotropic roles in protection against infections and inflammation, predominantly as a nutrient substrate for commensal microbial communities supporting their activity and improving the health of the host. It can however, provide a competitive advantage for certain pathogens. Fucose utilization appears to be important for the survival of *C.jejuni* NCTC11168 in the gut [14] and the pathogen exhibits chemotaxis and increased biofilm formation in the presence of fucose [16]. The correlation we see between LCN-2 and fucose here is a protective host immune response or against *C.jejuni* 81-176, since this strain lacks the *fuc* locus that regulates chemotaxis towards fucose and therefore the pathogen cannot use fucose to its benefit. Elevated urinary fucose excretion accompanied by increased LCN-2 levels were also observed in our murine model of *Cryptosporidium* infection [34]. We did not identify any ASVs belonging to the *Campylobacter* genus within our study. Interestingly, a previous study on a murine model of *C. jejuni* infection also did not report the presence of sequences assigned to *Campylobacter* [48]. In this study, the authors amplified the V35 region of the 16s rRNA gene, while the V4 region was chosen for our study. The lack of detection of *Campylobacter* even when qPCR indicates abundant shedding suggests that existing universal primers for amplification of 16S rRNA genes may have bias that prevents amplification of 16S rRNA genes from *Campylobacter* species.

The specific host nutrient and microbiome alterations that alter relevant disease outcomes offer another potentially fruitful area to better dissect and understand the mechanisms (and thus potentially effective approaches to intervene) for these clinical effects of *Campylobacter* infections. Thus we can now explore whether zinc or key repair nutrients might ameliorate these disease outcomes. For example, zinc can suppress expression of not only such host genes as the CFTR chloride channel that is responsible for secretory diarrhea with cholera and the *muc-2* gene, but also microbial genes such as the *aggR*-regulated virulence gene expression in enteroaggregative *E. coli* (EAEC) *in vitro* or *in vivo* [33], and homologous genes have been reported in several other enteric bacterial pathogens like *Vibrio cholerae*, *Shigella* and *Citrobacter,* and possibly *Campylobacter* [49, 50]. Effects of zinc on enteric infections are complex. For example, in addition to our findings herein regarding *C. jejuni* infection, we have also recently reported enhanced enteroaggregative *E. coli* infection in zinc deficient mice [33, 51]. However, the opposite is seen with *C. difficile* infection which is worse with added zinc, potentially via altering gut microbiota, toxin production and even the zinc binding host antimicrobial protein, calprotectin [52].

We conclude that zinc deficiency and antimicrobial treatment enable a robust weaned murine model of symptomatic enteric *Campylobacter* infection that exhibits its major clinical manifestation in humans of bloody diarrhea and growth failure, inflammatory histopathology and biomarker expression and fecal shedding that is sustained over at least 2 weeks. Others have explored the role of zinc and innate immune response in the pathogenesis of *Campylobacte*r infections. Interestingly, Davis *et al*. describes the requirement of a zinc transporter (znuA) for growth of *Campylobacter* in zinc-limiting media and in chickens. The same wildtype strain was used in this study (81-176) and was able to colonize in zinc deficient mice [53, 54]. Whether or not a znuA mutant could colonize in this mouse model is unknown. Whether zinc deficiency benefits *Campylobacter* (similar to our findings with Enteroaggregative *Escherichia coli* [33]) or alters normal microbiota in combination with antibiotics requires further investigation. Studies are currently underway to address these possible effects of zinc deficiency for both host and pathogen mechanisms. This may also help explain the requirement for antibiotic disruption of native flora prior to *Campylobacte*r challenge in our model. In contrast, with either standard or protein deficient diets (instead of zinc deficiency), we see modest enteropathy (as demonstrated by shedding and biomarkers), without overt diarrhea, thus potentially modeling ‘asymptomatic’ *Campylobacter* infections in children in developing areas worldwide.

In conclusion, these models now enable more rapid and clinically relevant vaccine testing and dissection of mechanisms that may help in development of novel approaches to effective interventions for this common global threat.

## Methods

### Animal husbandry

This study included the use of mice. This study was carried out in strict accordance with the recommendations in the Guide for the Care and Use of Laboratory Animals of the National Institutes of Health. The protocol was approved by the Committee on the Ethics of Animal Experiments of the University of Virginia (Protocol Number: 3315). All efforts were made to minimize suffering. This protocol was approved and is in accordance with the Institutional Animal Care and Use Committee policies of the University of Virginia. The University of Virginia is accredited by the Association for the Assessment and Accreditation of Laboratory Animal Care, International (AAALAC). Mice used in this study were male, 22 days old, C57BL/6 strain, and ordered from Jackson Laboratories (Bar Harbor, ME). Mice weighed approximately 11 grams on arrival and were co-housed in groups up to five animals per cage. The vivarium was kept at a temperature of between 68-74°F (20-23°C) with a 14 hour light and 10 hour dark cycle.

### Rodent Diet

Weaned mice (22 days old) were acclimated, fed a regular diet for 2-5 days, and then were fed either standard rodent ‘House Chow’ (HC), a protein source diet without zinc (dZD), or protein (2%) deficient (dPD) diet (Research Diets, Inc.). All diets were isocaloric and calories from fat, protein, and carbohydrates are as previously reported [35].

### *C. jejuni* Infection

Weaned, 3 week old male Jackson mice were acclimated for 3 days on standard rodent house chow (HC). Mice were then given HC, dZD or dPD described above for 14 days before challenge. Prior to infection, mice were given gentamicin (35mg/L), vancomycin (45mg/L), metronidazole (215mg/L), and colistin (850U/ml) in drinking water to disrupt resident microbiota as previously published [33, 55]. After three days on antibiotic water, mice were given untreated water without antibiotics for 1 day prior to challenge. *C. jejuni* 81-176 and an isogenic *kpsM* mutant were used [37]. Cultures were grown from glycerol stocks in Mueller Hinton broth in airtight canisters with ‘campy packs’ (Fisher Scientific). Each infected mouse received an inoculum ∼1×10^7^ *C. jejuni* in 100 µL of freshly prepared broth via oral gavage; controls received 100 µL of broth alone. Mice were maintained on specific diet and untreated filtered drinking water following infection.

Each experiment used N=8 mice per experimental group and mice were euthanized at either day 3 or day 14 post infection. We have previously published that we obtain both serum and tissue zinc deficiency following 14 days on dZD diet [33].

### Protein extraction from stool and tissue

After rapid dissection of the mouse intestines, cecal contents and stool were flash frozen in liquid nitrogen and stored in a -80C freezer. At time of assay, samples were lysed in RIPA buffer (20 mM Tris, pH 7.5, 150 mM NaCl, 1% Nonidet P-40, 0.5% sodium deoxycholate, 1 mM EDTA, 0.1% SDS) containing protease inhibitors cocktail (Roche) and phosphatase inhibitors (1 mM sodium orthovanadate, 5 mM sodium fluoride, 1 mM microcystin LR, and 5 mM beta-glycerophosphate). Tissue lysates were cleared by centrifugation, and the supernatant was used for total protein measurement (Pierce BCA), cytokine measurement by Luminex assay (BioRad), and specific ELISAs for lipocalin-2 (LCN2) and myeloperoxidase (MPO) as previously described [35].

### DNA extraction

DNA was isolated from cecal contents and fecal pellets using the QIAamp DNA stool mini kit (Qiagen) as previously described [56]. For enhancing the pathogen’s DNA extraction we made an improvement in the original protocol: a vigorous homogenization of the samples with 300 mg of 1.0 mm zirconia beads (BioSpec, Bartlesville, OK, USA) using a MiniBeadBeater (BioSpec, Bartlesville, OK, USA). After extraction, DNA was eluted in 200μl Elution Buffer and stored at −20°C.

### Real-time PCR for *C. jejuni* quantification

Stool and tissue DNA were analyzed for the *C. jejuni/coli cadF* gene to determine shedding of organism in stool or burden in the tissue. Primers for *cadF* were F, ‘CTGCTAAACCATAGAAATAAAATTTCTCAC’; R, ‘CTTTGAAGGTAATTTAGATATGGATAATCG’; Probe, ‘FAM-CATTTTGACGATTTTTGGCTTGA-MGB’ as published by Liu *et al*. [57]. Quantification of the infection was performed in a Bio-Rad CFX PCR Detection System by interpolating Ct values of each run with a standard curve of known amounts of *C. jejuni* DNA and transformed into number of organisms per milligram of sample. The master mix solution and primers were used as described elsewhere (3). Amplification consisted of 5 minutes at 95 °C, followed by 40 cycles of 15 seconds at 95 °C, 60 seconds at 58°C. PCR efficiency was >90% with reliable detection from 10^2^-10^10^ cfu as determined with a standard curve verified by plate counts.

### Intestinal morphometry and mucus staining

Ileal segments were fixed in 4% paraformaldehyde, embedded in paraffin, and stained with hematoxylin-eosin or PAS at the University of Virginia Histology Core.

### Biomarkers

Myeloperoxidase (MPO) and lipocalin-2 (LCN-2) were measured in stool and cecal contents according to manufacturer’s instructions (R&D Systems). Values were normalized to total protein as measured by BCA kit (Invitrogen) and presented as pg/mg protein.

### Statistical analysis

Data analyses were performed with GraphPad Prism 6 software (GraphPad Software). All statistical analyses were done from raw data with the use of analysis of variance, Student t tests, and Bonferroni post hoc analysis where applicable. Differences were considered significant at P < 0.05. Data are represented as means ± standard errors of the mean.

### ^1^H NMR spectroscopy based metabolic profiling

Urine samples were analysed by ^1^H nuclear magnetic resonance (NMR) spectroscopy. Each sample was prepared by combining 30 μl of urine with 30 μl of phosphate buffer (pH 7.4, 100% D_2_O) containing 1mM of the internal standard, 3-trimethylsilyl-1-[2,2,3,3-^2^H4] propionate (TSP). Samples were then vortexed and spun (10,000 g) for 10 minutes at 4 °C before transfer to 1.7mm NMR tube. Spectroscopic analysis was performed at 300K on a 600 MHz Bruker NMR spectrometer equipped with a BBI probe. Standard one-dimensional spectra of the urine samples were acquired with saturation of the water resonance, using a standard pulse sequence. For each sample, 4 dummy scans were followed by 64 scans collected in 64K time domain points and with a spectral window set to 20 ppm. A relaxation delay of 4 s, a mixing time of 10 ms, an acquisition time of 2.73 s and 0.3 Hz line broadening was used. Spectra were referenced to the TSP resonance at δ 0.0. ^1^H NMR spectra (δ -0.5 – 10) were digitized into consecutive integrated spectral regions (∼ 20,000) of equal width (0.00055 ppm). Spectral regions corresponding to TSP (δ -0.5 – 0.5), water (δ 4.5 – 4.8) and urea (δ 5.6 – 6.1), were removed. The resulting spectra were then normalized to unit area. Multivariate statistical modeling was performed using in-house scripts, In MATLAB (R2016a). This included principal components analysis (PCA) using Pareto scaling and orthogonal projections to latent structures-discriminant analysis (OPLS-DA) using data mean centering. OPLS-DA models were built to facilitate data interpretation. ^1^H NMR spectroscopic profiles were used as the descriptor matrix and class membership (e.g., uninfected zinc deficient mice and *Campylobacter* infected zinc deficient mice) was used as the response variable. The predictive performance (Q^2^Y) of the models was calculated with the use of a 7-fod cross-validation method, and model validity was defined upon permutation testing (1000 permutations). Significant metabolites were identified and extracted from valid pair-wise OPLS-DA models and summarized in heat maps.

### 16S rRNA gene sequencing and analysis

16S libraries were pooled and sequenced according to the protocol developed by Kozich *et al*., and the detailed version of the specific protol used is provided in the supplemental text. Briefly, the V4 region of the 16S rRNA gene was amplified and sequenced using the MiSeq Reagent Kit v2 that produces up to 25 million reads of 2 x 250 bp per run at the Genomics Core Facility at UVA. Reads were assigned to samples using Illumina BaseSpace demultiplexing. From these reads, analysis was performed using the DADA2 package (benjjneb.github.io/dada2, version 1.4.0) [58]. The first 10 bases and last 10 bases were trimmed from each 250 base forward read due to low quality scores, resulting in 230 base sequences. Similarly, the first 10 bases and the last 90 bases were trimmed from reverse reads, resulting in 150 base sequences. Reads were then filtered according to 3 criteria: presence of no ambiguous ‘N’ bases in the read, no base with quality scores below 2 in the read, and having a maximum of 2 expected errors in the read. Filtered reads were then dereplicated with the derepFastq function, resulting in unique sequences each with a corresponding abundance and consensus quality profile. The DADA2 sample inference function was applied to dereplicated sequences with joint inference of sample composition and error-rate parameter estimation (e.g. selfConsist = TRUE). Chimeras were then removed using the removeBimeraDenovo function, then taxonomy was assigned to each sequence using the Ribosomal Database Project’s naïve bayes classifier [59] with RDP training set 14.

Amplicon sequence variant (ASV) counts for each sample, with accompanying taxonomy assignments, were further analysed using the phyloseq [60] package (version 1.20) in the R programming language (version 3.4.0). Differentially abundant ASVs were determined using the DESeq2 [61] package (version 1.16.1). Prior to all analyses, sample counts were rarified to control for uneven sequencing depth across samples. For each sample, 5000 reads were randomly chosen using the rarefy_even_depth function in phyloseq. These rarefied counts were used for all downstream analyses. Alpha diversity metrics were calculated using the plot_richness function in phyloseq. A three-way ANOVA was used to compare richness over time, between diets, and between infection statuses, with multiple testing correction via Tukey’s Honestly Significant Difference (HSD) method as implemented in the TukeyHSD function. Comparisons with a multiple testing corrected p value of less than 0.05 were considered significantly different. The same procedure was used to compare evenness across sample groups. Bray-Curtis dissimilarity calculations and non-metric multidimensional scaling (k=2) were performed using the plot_ordination function in phyloseq. Differential abundance testing was performed between time-matched infected and uninfected mice within each diet. Prior to differential abundance testing, ASVs with fewer than 10 counts in any sample or fewer than 20 counts across the entire dataset were removed. Within DESeq2, Wald significance tests were used with parametric dispersion fitting. ASVs with a p value of less than 0.05 were considered significantly differentially abundant. The DESeq2 design was chosen to compare groups across all three-group categories (infection status, diet, and time), and the results were extracted specifically for comparisons between infected and uninfected groups with matched diets at the same time points. ASVs that were differentially abundant in any of these infected vs. uninfected comparisons are included in the heatmap in Fig 9. DADA2 processing scripts, processed data as a phyloseq object, and scripts to recreate all microbiome analyses can be found at https://github.com/gregmedlock/campy_murine_diets. Raw sequence data are currently being submitted to NCBI SRA.

## Author contributions

NG: Data Curation, Formal analysis, Investigation, Methodology, Software, Visualization, Writing - Original Draft Preparation, Writing – Review & Editing

GLM: Data Curation, Formal analysis, Investigation, Methodology, Software, Visualization, Writing - Original Draft Preparation, Writing – Review & Editing

DTB: Conceptualization, Data Curation, Formal Analysis, Investigation, Methodology, Validation, Visualization, Writing - Original Draft Preparation, Writing – Review & Editing

PHQSM: Investigation, Writing – Review & Editing

SEL: Investigation, Writing – Review & Editing

GLK: Conceptualization, Resources, Writing – Review & Editing

KT: Investigation, Resources

GP: Resources

JRS: Conceptualization, Funding Acquisition, Project Administration, Supervision, Resources, Writing – Review & Editing

RLG: Conceptualization, Funding Acquisition, Project Administration, Supervision, Resources, Writing – Review & Editing

## Supporting Information

**S1 Fig.** *C. jejuni* grown on selection agar (Hardy Diagnostics, Santa Maria, CA) from a single fecal pellet diluted in PBS 10^-4^ one day post infection. Image shown is representative of findings from 8 infected mice.

**S2 Fig.** ‘Swiss roll’ PAS stained histology of colon from house chow fed (**Panel A**) and zinc deficient diet fed (**Panel B**) mice 3 days following infection with *Campylobacter jejuni*. Notable changes in the zinc deficient mice were increased lamina propria cellular infiltrate and striking lumenal mucus discharge with only minor mucus discharge seen in house chow fed mice.

**S3 Fig. OPLS-DA coefficients plot comparing the urinary metabolic profiles of HC-fed mice and dZD-fed mice.** (HC: n= 8, dZD n=8, (Q^2^Y=0.45, P = 0.005) on day 0. Positive peaks correspond to metabolites being excreted in higher amounts by the dZD-fed mice and negative peaks indicate metabolites excreted in lower amounts as, compared to the HC-fed mice. Abbreviations: 2-OG, 2-oxoglutarate; 2-OIV, 2-oxoisovalerate; 2-OIC, 2-oxoisocaproate; 2-MOV, 3-methyl-2-oxovalerate; 2-PY, N-methyl-2-pyridone-5-carboxamide; 4-HPA, 4-hydroxyphenylacetate; BG, butyrylglycine; DMG, dimethylglycine; HG, hexanoylglycine; IV, isovalerate; NMNA, N-methyl-nicotinic acid; NMND, N-methylnicotinamide; TMA, trimethylamine.

**S1 File MiSeq Wet Lab SOP**

## References

1. Scallan E, Hoekstra RM, Mahon BE, Jones TF, Griffin PM. An assessment of the human health impact of seven leading foodborne pathogens in the United States using disability adjusted life years. Epidemiology and infection. 2015;143(13):2795–804. Epub 2015/01/31. doi: 10.1017/S0950268814003185. PubMed PMID: 25633631.

2. Kirk MD, Pires SM, Black RE, Caipo M, Crump JA, Devleesschauwer B, et al. World Health Organization Estimates of the Global and Regional Disease Burden of 22 Foodborne Bacterial, Protozoal, and Viral Diseases, 2010: A Data Synthesis. PLoS medicine. 2015;12(12):e1001921. Epub 2015/12/04. doi: 10.1371/journal.pmed.1001921. PubMed PMID: 26633831; PubMed Central PMCID: PMC4668831.

3. Kotloff KL, Nataro JP, Blackwelder WC, Nasrin D, Farag TH, Panchalingam S, et al. Burden and aetiology of diarrhoeal disease in infants and young children in developing countries (the Global Enteric Multicenter Study, GEMS): a prospective, case-control study. Lancet. 2013;382 (9888):209-22. Epub 2013/05/18. doi: 10.1016/S0140-6736(13)60844-2. PubMed PMID: 23680352.

4. Platts-Mills JA, Babji S, Bodhidatta L, Gratz J, Haque R, Havt A, et al. Pathogen-specific burdens of community diarrhoea in developing countries: a multisite birth cohort study (MAL-ED). The Lancet Global health. 2015;3(9):e564–75. Epub 2015/07/24. doi: 10.1016/S2214-109X(15)00151-5. PubMed PMID: 26202075.

5. Amour C, Gratz J, Mduma E, Svensen E, Rogawski ET, McGrath M, et al. Epidemiology and Impact of Campylobacter Infection in Children in 8 Low-Resource Settings: Results From the MAL-ED Study. Clin Infect Dis. 2016;63(9):1171–9. Epub 2016/08/10. doi: 10.1093/cid/ciw542. PubMed PMID: 27501842; PubMed Central PMCID: PMC5064165.

6. Allos B, Lastovica A. Tropical infectious diseases : principles, pathogens and practice. Third edition. ed. Edinburgh: Saunders/Elsevier; 2011.

7. Blaser MJ, Parsons RB, Wang WL. Acute colitis caused by Campylobacter fetus ss. jejuni. Gastroenterology. 1980;78(3):448–53. Epub 1980/03/01. PubMed PMID: 7351284.

8. Blaser MJ, Wells JG, Feldman RA, Pollard RA, Allen JR. Campylobacter enteritis in the United States. A multicenter study. Ann Intern Med. 1983;98(3):360–5. Epub 1983/03/01. PubMed PMID: 6830079.

9. Williams MD, Schorling JB, Barrett LJ, Dudley SM, Orgel I, Koch WC, et al. Early treatment of Campylobacter jejuni enteritis. Antimicrob Agents Chemother. 1989;33(2):248–50. Epub 1989/02/01. PubMed PMID: 2818711; PubMed Central PMCID: PMC171468.

10. Stahl M, Frirdich E, Vermeulen J, Badayeva Y, Li X, Vallance BA, et al. The Helical Shape of Campylobacter jejuni Promotes In Vivo Pathogenesis by Aiding Transit through Intestinal Mucus and Colonization of Crypts. Infect Immun. 2016;84(12):3399–407. doi: 10.1128/IAI.00751-16. PubMed PMID: 27647867; PubMed Central PMCID: PMCPMC5116718.

11. Stahl M, Vallance BA. Insights into Campylobacter jejuni colonization of the mammalian intestinal tract using a novel mouse model of infection. Gut Microbes. 2015;6(2):143–8. doi: 10.1080/19490976.2015.1016691. PubMed PMID: 25831043; PubMed Central PMCID: PMCPMC4615362.

12. Stahl M, Ries J, Vermeulen J, Yang H, Sham HP, Crowley SM, et al. A novel mouse model of Campylobacter jejuni gastroenteritis reveals key pro-inflammatory and tissue protective roles for Toll-like receptor signaling during infection. PLoS Pathog. 2014;10(7):e1004264. doi: 10.1371/journal.ppat.1004264. PubMed PMID: 25033044; PubMed Central PMCID: PMCPMC4102570.

13. Stahl M, Butcher J, Stintzi A. Nutrient acquisition and metabolism by Campylobacter jejuni. Front Cell Infect Microbiol. 2012;2:5. doi: 10.3389/fcimb.2012.00005. PubMed PMID: 22919597; PubMed Central PMCID: PMCPMC3417520.

14. Stahl M, Friis LM, Nothaft H, Liu X, Li J, Szymanski CM, et al. L-fucose utilization provides Campylobacter jejuni with a competitive advantage. Proceedings of the National Academy of Sciences of the United States of America. 2011;108(17):7194–9. doi: 10.1073/pnas.1014125108. PubMed PMID: 21482772; PubMed Central PMCID: PMCPMC3084102.

15. Weingarten RA, Grimes JL, Olson JW. Role of Campylobacter jejuni respiratory oxidases and reductases in host colonization. Applied and environmental microbiology. 2008;74(5):1367–75. Epub 2008/01/15. doi: 10.1128/AEM.02261-07. PubMed PMID: 18192421; PubMed Central PMCID: PMC2258625.

16. Dwivedi R, Nothaft H, Garber J, Xin Kin L, Stahl M, Flint A, et al. L-fucose influences chemotaxis and biofilm formation in Campylobacter jejuni. Mol Microbiol. 2016;101(4):575–89. doi: 10.1111/mmi.13409. PubMed PMID: 27145048.

17. Lopez GU, Kitajima M, Sherchan SP, Sexton JD, Sifuentes LY, Gerba CP, et al. Impact of disinfectant wipes on the risk of Campylobacter jejuni infection during raw chicken preparation in domestic kitchens. J Appl Microbiol. 2015;119(1):245–52. Epub 2015/05/06. doi: 10.1111/jam.12834. PubMed PMID: 25939813.

18. Sahin O, Yaeger M, Wu Z, Zhang Q. Campylobacter-Associated Diseases in Animals. Annu Rev Anim Biosci. 2016. PubMed PMID: Medline:27860495.

19. Liu J, Kabir F, Manneh J, Lertsethtakarn P, Begum S, Gratz J, et al. Development and assessment of molecular diagnostic tests for 15 enteropathogens causing childhood diarrhoea: a multicentre study. Lancet Infect Dis. 2014;14(8):716–24. Epub 2014/07/16. doi: 10.1016/S1473-3099(14)70808-4. PubMed PMID: 25022434.

20. Riddle MS, Guerry P. Status of vaccine research and development for Campylobacter jejuni. Vaccine. 2016;34(26):2903–6. PubMed PMID: Medline:26973064.

21. Marder EP, Cieslak PR, Cronquist AB, Dunn J, Lathrop S, Rabatsky-Ehr T, et al. Incidence and Trends of Infections with Pathogens Transmitted Commonly Through Food and the Effect of Increasing Use of Culture-Independent Diagnostic Tests on Surveillance - Foodborne Diseases Active Surveillance Network, 10 US Sites, 2013-2016. Mmwr-Morbid Mortal W. 2017;66(15):397-403. PubMed PMID: WOS:000399550500001.

22. Babakhani FK, Bradley GA, Joens LA. Newborn piglet model for campylobacteriosis. Infect Immun. 1993;61(8):3466–75. Epub 1993/08/01. PubMed PMID: 8335377; PubMed Central PMCID: PMC281024.

23. Fox JG, Ackerman JI, Taylor N, Claps M, Murphy JC. Campylobacter jejuni infection in the ferret: an animal model of human campylobacteriosis. Am J Vet Res. 1987;48(1):85–90. Epub 1987/01/01. PubMed PMID: 3826848.

24. Askoura M, Stintzi A. Using Galleria mellonella as an Infection Model for Campylobacter jejuni Pathogenesis. Methods in molecular biology. 2017;1512:163–9. Epub 2016/11/26. doi: 10.1007/978-1-4939-6536-6_14. PubMed PMID: 27885606.

25. Davis L, DiRita V. Experimental chick colonization by Campylobacter jejuni. Current protocols in microbiology. 2008;Chapter 8:Unit 8A 3. Epub 2008/11/20. doi: 10.1002/9780471729259.mc08a03s11. PubMed PMID: 19016444; PubMed Central PMCID: PMC5147583.

26. Johnson JG, DiRita VJ. Generation and Screening of an Insertion Sequencing-Compatible Mutant Library of Campylobacter jejuni. Methods in molecular biology. 2017;1512:257–72. Epub 2016/11/26. doi: 10.1007/978-1-4939-6536-6_21. PubMed PMID: 27885613.

27. Bereswill S, Fischer A, Plickert R, Haag LM, Otto B, Kuhl AA, et al. Novel murine infection models provide deep insights into the “menage a trois” of Campylobacter jejuni, microbiota and host innate immunity. PLoS One. 2011;6(6):e20953. Epub 2011/06/24. doi: 10.1371/journal.pone.0020953. PubMed PMID: 21698299; PubMed Central PMCID: PMC3115961.

28. Iizumi T, Taniguchi T, Yamazaki W, Vilmen G, Alekseyenko AV, Gao Z, et al. Effect of antibiotic pre-treatment and pathogen challenge on the intestinal microbiota in mice. Gut pathogens. 2016;8:60. Epub 2016/11/29. doi: 10.1186/s13099-016-0143-z. PubMed PMID: 27891184; PubMed Central PMCID: PMC5116224.

29. Stahl M, Graef FA, Vallance BA. Mouse Models for Campylobacter jejuni Colonization and Infection. Methods in molecular biology. 2017;1512:171–88. Epub 2016/11/26. doi: 10.1007/978-1-4939-6536-6_15. PubMed PMID: 27885607.

30. Stanfield JT, McCardell BA, Madden JM. Campylobacter diarrhea in an adult mouse model. Microb Pathog. 1987;3(3):155–65. PubMed PMID: 3504219.

31. McCardell BA, Madden JM, Stanfield JT. Effect of iron concentration on toxin production in Campylobacter jejuni and Campylobacter coli. Can J Microbiol. 1986;32(5):395–401. PubMed PMID: 3719459.

32. McCardell BA, Madden JM, Stanfield JT. A mouse model for the measurement of virulence of species of Campylobacter. J Infect Dis. 1986;153(1):177. PubMed PMID: 3941285.

33. Bolick DT, Kolling GL, Moore JH, 2nd, de Oliveira LA, Tung K, Philipson C, et al. Zinc deficiency alters host response and pathogen virulence in a mouse model of enteroaggregative escherichia coli-induced diarrhea. Gut Microbes. 2014;5(5):618–27. Epub 2014/12/09. doi: 10.4161/19490976.2014.969642. PubMed PMID: 25483331.

34. Bolick DT, Mayneris-Perxachs J, Medlock GL, Kolling GL, Papin J, Swann JR, et al. Increased urinary trimethylamine N-oxide (TMAO) following Cryptosporidium infection and protein malnutrition independent of microbiome effects. J Infect Dis. 2017. Epub 2017/05/19. doi: 10.1093/infdis/jix234. PubMed PMID: 28520899.

35. Mayneris-Perxachs J, Bolick DT, Leng J, Medlock GL, Kolling GL, Papin JA, et al. Protein- and zinc-deficient diets modulate the murine microbiome and metabolic phenotype. Am J Clin Nutr. 2016. PubMed PMID: Medline:27733402.

36. Prata MMG HA, Bolick DT, Pinkerton R, Lima AMM, Guerrant RL. Comparisons between myeloperoxidase, lactoferrin, calprotectin and lipocalin-2, as fecal biomarkers of intestinal inflammation in malnourished children. J Transl Sci. 2016;2(2):134–9. doi: 10.15761/JTS.1000130.

37. Bacon DJ, Szymanski CM, Burr DH, Silver RP, Alm RA, Guerry P. A phase-variable capsule is involved in virulence of Campylobacter jejuni 81-176. Mol Microbiol. 2001;40(3):769–77. Epub 2001/05/22. PubMed PMID: 11359581.

38. Winter SE, Lopez CA, Baumler AJ. The dynamics of gut-associated microbial communities during inflammation. EMBO Rep. 2013;14(4):319–27. doi: 10.1038/embor.2013.27. PubMed PMID: 23478337; PubMed Central PMCID: PMCPMC3615657.

39. Maue AC, Mohawk KL, Giles DK, Poly F, Ewing CP, Jiao Y, et al. The polysaccharide capsule of Campylobacter jejuni modulates the host immune response. Infect Immun. 2013;81(3):665–72. Epub 2012/12/20. doi: 10.1128/IAI.01008-12. PubMed PMID: 23250948; PubMed Central PMCID: PMC3584872.

40. St Charles JL, Bell JA, Gadsden BJ, Malik A, Cooke H, Van de Grift LK, et al. Guillain Barre Syndrome is induced in Non-Obese Diabetic (NOD) mice following Campylobacter jejuni infection and is exacerbated by antibiotics. Journal of autoimmunity. 2016. Epub 2016/12/13. doi: 10.1016/j.jaut.2016.09.003. PubMed PMID: 27939129.

41. Kosek M, Haque R, Lima A, Babji S, Shrestha S, Qureshi S, et al. Fecal markers of intestinal inflammation and permeability associated with the subsequent acquisition of linear growth deficits in infants. Am J Trop Med Hyg. 2013;88(2):390–6. Epub 2012/11/28. doi: 10.4269/ajtmh.2012.12-0549. PubMed PMID: 23185075; PubMed Central PMCID: PMC3583335.

42. Guerrant RL, Leite AM, Pinkerton R, Medeiros PH, Cavalcante PA, DeBoer M, et al. Biomarkers of Environmental Enteropathy, Inflammation, Stunting, and Impaired Growth in Children in Northeast Brazil. PLoS One. 2016;11(9):e0158772. doi: 10.1371/journal.pone.0158772. PubMed PMID: 27690129; PubMed Central PMCID: PMCPMC5045163.

43. Kraus D, Yang Q, Kong D, Banks AS, Zhang L, Rodgers JT, et al. Nicotinamide N-methyltransferase knockdown protects against diet-induced obesity. Nature. 2014;508(7495):258-62. doi: 10.1038/nature13198. PubMed PMID: 24717514; PubMed Central PMCID: PMCPMC4107212.

44. Mayneris-Perxachs J, Bolick DT, Leng J, Medlock GL, Kolling GL, Papin JA, et al. Protein- and zinc-deficient diets modulate the murine microbiome and metabolic phenotype. Am J Clin Nutr. 2016;104(5):1253–62. doi: 10.3945/ajcn.116.131797. PubMed PMID: 27733402; PubMed Central PMCID: PMCPMC5081716.

45. Brand IA, Kleineke J. Intracellular zinc movement and its effect on the carbohydrate metabolism of isolated rat hepatocytes. The Journal of biological chemistry. 1996;271(4):1941–9. PubMed PMID: 8567642.

46. Dawson LF, Donahue EH, Cartman ST, Barton RH, Bundy J, McNerney R, et al. The analysis of para-cresol production and tolerance in Clostridium difficile 027 and 012 strains. BMC Microbiol. 2011;11:86. doi: 10.1186/1471-2180-11-86. PubMed PMID: 21527013; PubMed Central PMCID: PMCPMC3102038.

47. Nothaft H, Perez-Munoz ME, Gouveia GJ, Duar RM, Wanford JJ, Lango-Scholey L, et al. Co-administration of the Campylobacter jejuni N-glycan based vaccine with probiotics improves vaccine performance in broiler chickens. Appl Environ Microbiol. 2017. doi: 10.1128/AEM.01523-17. PubMed PMID: 28939610; PubMed Central PMCID: PMCPMC5691412.

48. O’Loughlin JL, Samuelson DR, Braundmeier-Fleming AG, White BA, Haldorson GJ, Stone JB, et al. The Intestinal Microbiota Influences Campylobacter jejuni Colonization and Extraintestinal Dissemination in Mice. Appl Environ Microbiol. 2015;81(14):4642–50. doi: 10.1128/AEM.00281-15. PubMed PMID: 25934624; PubMed Central PMCID: PMCPMC4551207.

49. Santiago AE, Yan MB, Tran M, Wright N, Luzader DH, Kendall MM, et al. A large family of anti-activators accompanying XylS/AraC family regulatory proteins. Mol Microbiol. 2016;101(2):314–32. Epub 2016/04/03. doi: 10.1111/mmi.13392. PubMed PMID: 27038276; PubMed Central PMCID: PMC4983702.

50. Flom Gary A. Pathogenic mechanisms of Campylobacter jejuni : regulation of Campylobacter invasion antigen B [PDF file.]. 2004. Available from: http://www.dissertations.wsu.edu/thesis/Spring2004/G_Flom_050704.pdf.

51. Medeiros P, Bolick DT, Roche JK, Noronha F, Pinheiro C, Kolling GL, et al. The micronutrient zinc inhibits EAEC strain 042 adherence, biofilm formation, virulence gene expression, and epithelial cytokine responses benefiting the infected host. Virulence. 2013;4(7). Epub 2013/08/21. PubMed PMID: 23958904.

52. Zackular JP, Moore JL, Jordan AT, Juttukonda LJ, Noto MJ, Nicholson MR, et al. Dietary zinc alters the microbiota and decreases resistance to Clostridium difficile infection. Nat Med. 2016;22(11):1330–4. Epub 2016/11/01. doi: 10.1038/nm.4174. PubMed PMID: 27668938; PubMed Central PMCID: PMC5101143.

53. Gielda LM, DiRita VJ. Zinc competition among the intestinal microbiota. mBio. 2012;3(4):e00171–12. Epub 2012/08/02. doi: 10.1128/mBio.00171-12. PubMed PMID: 22851657; PubMed Central PMCID: PMC3419517.

54. Davis LM, Kakuda T, DiRita VJ. A Campylobacter jejuni znuA orthologue is essential for growth in low-zinc environments and chick colonization. J Bacteriol. 2009;191(5):1631–40. Epub 2008/12/24. doi: 10.1128/JB.01394-08. PubMed PMID: 19103921; PubMed Central PMCID: PMC2648198.

55. Chen X, Katchar K, Goldsmith JD, Nanthakumar N, Cheknis A, Gerding DN, et al. A mouse model of Clostridium difficile-associated disease. Gastroenterology. 2008;135(6):1984–92. Epub 2008/10/14. doi: 10.1053/j.gastro.2008.09.002. PubMed PMID: 18848941.

56. Liu J, Bolick DT, Kolling GL, Fu Z, Guerrant RL. Protein Malnutrition Impairs Intestinal Epithelial Cell Turnover, a Potential Mechanism of Increased Cryptosporidiosis in a Murine Model. Infect Immun. 2016;84(12):3542–9. doi: 10.1128/IAI.00705-16. PubMed PMID: 27736783; PubMed Central PMCID: PMCPMC5116730.

57. Liu J, Gratz J, Amour C, Kibiki G, Becker S, Janaki L, et al. A laboratory-developed TaqMan Array Card for simultaneous detection of 19 enteropathogens. J Clin Microbiol. 2013;51(2):472–80. Epub 2012/11/24. doi: 10.1128/JCM.02658-12. PubMed PMID: 23175269; PubMed Central PMCID: PMC3553916.

58. Callahan BJ, McMurdie PJ, Rosen MJ, Han AW, Johnson AJ, Holmes SP. DADA2: High-resolution sample inference from Illumina amplicon data. Nature methods. 2016;13(7):581–3. Epub 2016/05/24. doi: 10.1038/nmeth.3869. PubMed PMID: 27214047; PubMed Central PMCID: PMC4927377.

59. Wang Q, Garrity GM, Tiedje JM, Cole JR. Naive Bayesian classifier for rapid assignment of rRNA sequences into the new bacterial taxonomy. Applied and environmental microbiology. 2007;73(16):5261–7. Epub 2007/06/26. doi: 10.1128/AEM.00062-07. PubMed PMID: 17586664; PubMed Central PMCID: PMC1950982.

60. McMurdie PJ, Holmes S. phyloseq: an R package for reproducible interactive analysis and graphics of microbiome census data. PLoS One. 2013;8(4):e61217. Epub 2013/05/01. doi: 10.1371/journal.pone.0061217. PubMed PMID: 23630581; PubMed Central PMCID: PMC3632530.

61. Love MI, Huber W, Anders S. Moderated estimation of fold change and dispersion for RNA-seq data with DESeq2. Genome biology. 2014;15(12):550. Epub 2014/12/18. doi: 10.1186/s13059-014-0550-8. PubMed PMID: 25516281; PubMed Central PMCID: PMC4302049.

