## Supplementary figures and images for "A novel mouse model of *Campylobacter jejuni* enteropathy and diarrhea"

### Supplementary Materials

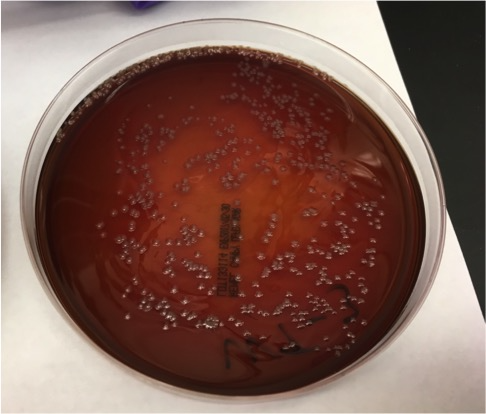

### Supplementary Materials

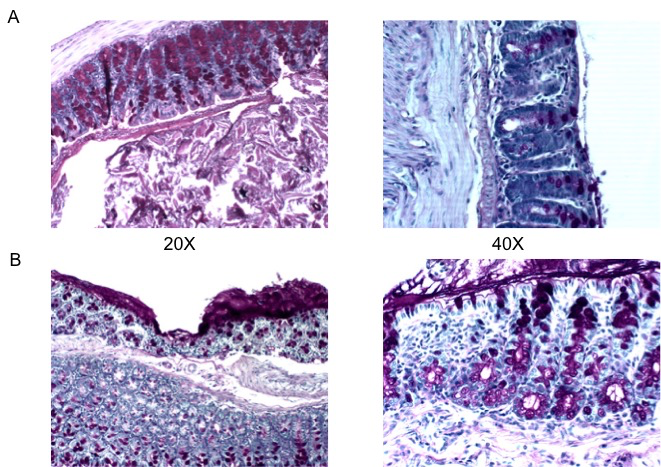

### Supplementary Materials

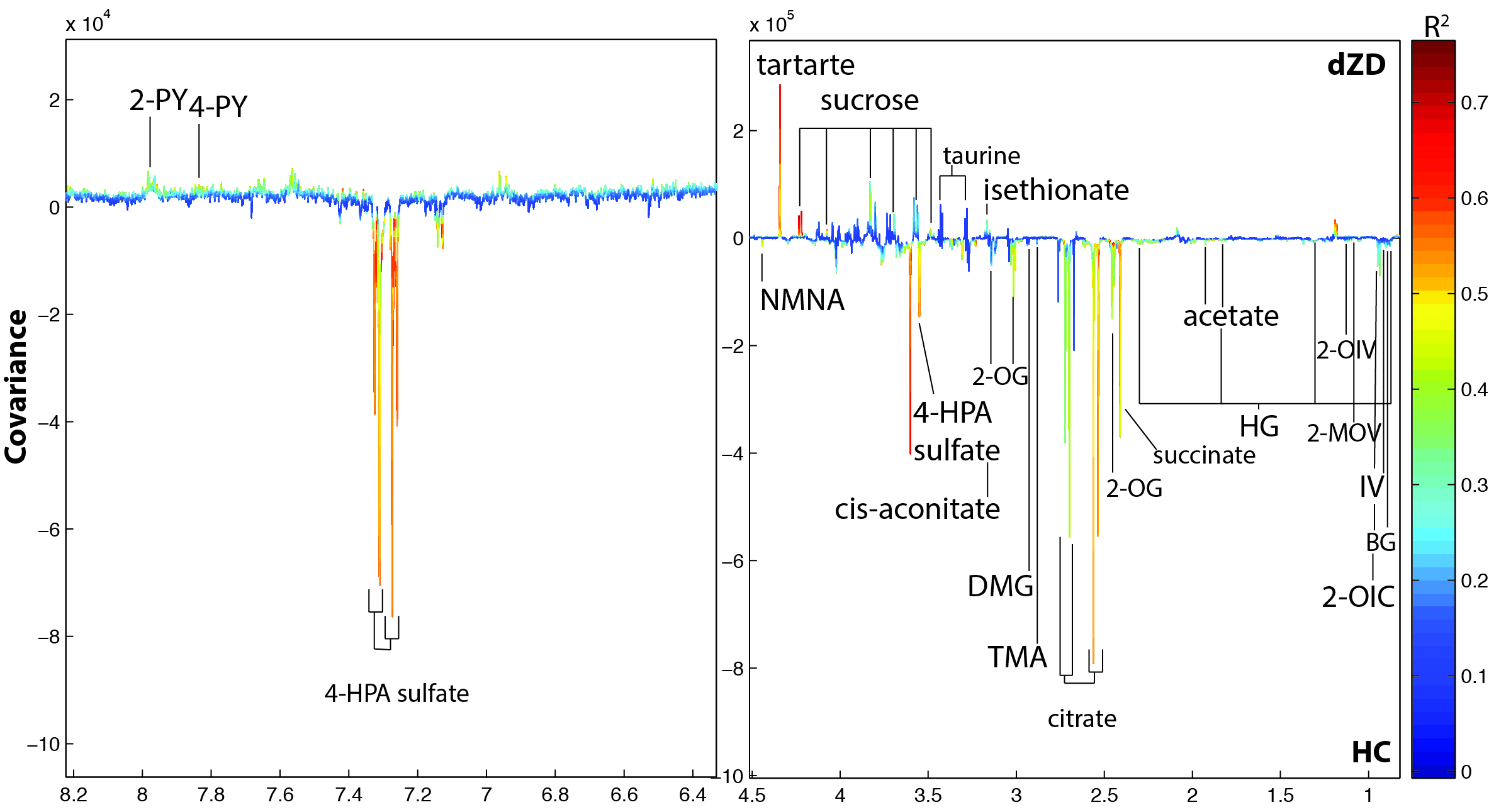
