## Supplementary Materials for "A novel mouse model of *Campylobacter jejuni* enteropathy and diarrhea"

**MiSeq Wet Lab SOP**

**Detailed Title:** | 16S rRNA Sequencing with the Illumina MiSeq: Library Generation, QC, & Sequencing
**Authors:** | James Kozich, Patrick Schloss, Niel Baxter, & Matt Jenior
**Original Date:** | 25-March-2013
**Version:** | 4.0
**Last Updated:** | 9-March-2015
**Updated By:** | Niel Baxter

**1.0) Introduction and Workflow**

***1.1) Introduction***

- The Purpose of this protocol is to define the steps for the preparation and sequencing of 16S rRNA gene sequence libraries using the Illumina MiSeq sequencing platform, as described in the paper [Development of a dual-index sequencing strategy and curation pipeline for analyzing amplicon sequence data on the MiSeq Illumina sequencing platform](http://www.ncbi.nlm.nih.gov/pubmed/23793624) by Kozich et al.
- The Illumina MiSeq Personal Sequencer can produce 2 x 250bp (or 2 x 300bp with v3 chemistry) paired-end reads and up to 8.5 Gb of data in a single run. Dual indexing of library samples allows up to 384 samples to be run simultaneously. The instrument is capable of producing in excess of 24 million reads. However for low diversity runs about 12 million reads can be expected. A wide range of applications is possible including 16S analysis, metagenomics, genome sequencing, transcriptomics, and RNA sequencing.
- Our lab typically sequences the V4 region of the 16S rRNA gene. Its short length (~250bp) allows for fully overlapping forward and reverse reads, which, in combination with our [curaton pipeline](http://www.mothur.org/wiki/MiSeq_SOP), results in the lowest error rates. We also prefer to use the older v2 MiSeq cartridges, as the newer v3 chemistry consistently results in a higher error rates in our hands.
- There are several steps in preparing samples for sequencing on the MiSeq. Broadly, these include library generation and indexing, quality control, normalization and pooling, quantification, sequencing, run quality assessment, and data export. An overview of each step and more detailed protocols are below.

***1.2) 16S Prep Workflow***

This section is an overview of the steps involved in library preparation.

For a more detailed desctiption of the methods, see Section 5 below.

1. Extracted DNA should be arrayed in 96 well plate format, preferably with two wells on each plate open for controls.
2. Samples are PCR amplified with Schloss lab indices. Each plate should contain a negative control (water) and a positive control (mock community)
3. A subset of 12-24 samples from each plate undergoes electrophoresis on a 1% agarose gel to ensure amplification proceeded normally.
4. Library clean up and normalization is performed using the Invitrogen [SequalPrep](http://products.invitrogen.com/ivgn/product/A1051001) Plate Normalization Kit.
5. Samples from each plate are pooled into single wells (i.e. 1 well/plate).
6. (Optional) To assess the quality of the library, the investigator may choose to perform a Bioanalyzer trace using the[Agilent Technologies HS DNAkit](http://www.genomics.agilent.com/CollectionSubpage.aspx?PageType=Product&SubPageType=ProductDetail&PageID=1635) cat# 5067-4626.
7. (Optional) If the post-PCR gel or the Bioanalyzer trace suggests contaminant DNA from leftover indices/primer-dimer, an additional gel purification of the pooled plates is recommended. This often improves the quality of the sequencing run.
8. Each pooled plate is quantified using a KAPA Biosystems [qPCR kit](http://www.kapabiosystems.com/products/name/kapa-library-quant-kits) cat# KK4824.
9. Plates are pooled to equal concentration into a single well (i.e. 1 well per run)
10. The pooled library enters the Sequencing Workflow.

***1.3) Sequencing Workflow***

This section is an overview of the steps involved in initiating a sequencing run.

For a more detailed desctiption of the mehods, see Section 5 below.

1. A Sample Plate is created for each plate using Illumina Experiment Manager. Sample Plates are then used to create a Sample Sheet. This sheet serves as the set of run parameters and indexing scheme used by the MiSeq for the run. The sample sheet is then transferred to the MiSeq via flash drive.
2. The reagent cartridge is thawed in a water bath per the [MiSeq System Guide](https://support.illumina.com/content/dam/illumina-support/documents/documentation/system_documentation/miseq/miseq-system-guide-15027617-01.pdf).
3. Unless otherwise specified, dilution and loading will follow the steps outlined in the document: [Preparing DNA Libraries for Sequencing on the MiSeq](https://support.illumina.com/content/dam/illumina-support/documents/documentation/system_documentation/miseq/preparing-libraries-for-sequencing-on-miseq-15039740-d.pdf)
   a. Pooled library and PhiX control are denatured and diluted.
   b. Diluted library and PhiX are pooled (5-10% PhiX, 90-95% Library).
   c. Sequencing pimers and library/PhiX are loaded into the reagent cartridge.
   d. MiSeq flow cell is washed
4. The sample sheet, flow cell, reagent cartridge, PR2 bottle, and an empty waste bottle are loaded onto the MiSeq, and the run is initiated. A 500 cycle run takes approx. 44 hours.
5. The run is monitored using Illumina Sequence Analysis Viewer.
6. Upon completions of the run, fastq files are transfered to the Schloss Lab NAS drive.
7. A post run wash is performed, followed by a standby wash if the machine will be idle for a week or more.

**2.0) Safety and Waste Disposal**

- The Schloss Lab Chemical Hygiene Plan should be followed at all times.
- Standard PPE (nitrile gloves, safety glasses, and lab coat) should be used at all times.
- Each reagent cartridge contains a small amount of formamide and must be disposed of in an appropriate container following the run. Liquid waste from a run must also be disposed of as hazardous due to the formamide content.

**3.0) Consumables**

**4.0) Run Costs**

| **For 384 sample run** | **PCR and Indexing** | **Cleanup & Normalization** | **Library QC** | **Sequencing** | **Totals** |
| --- | --- | --- | --- | --- | --- |
| 16S Reagents | $588 | $265 | $138 | $941 | $1932 |
| 16S Man hours | 4 | 3 | 4 | 4 | 15 |

**5.0) Detailed Method(s)**

***5.1) Published Protocols***

- The following methods and references are used in the workflows above.
  - [MiSeq System Guide](https://support.illumina.com/content/dam/illumina-support/documents/documentation/system_documentation/miseq/miseq-system-guide-15027617-01.pdf)
  - [Preparing DNA Libraries for Sequencing on the MiSeq](https://support.illumina.com/content/dam/illumina-support/documents/documentation/system_documentation/miseq/preparing-libraries-for-sequencing-on-miseq-15039740-d.pdf)
  - Kapa Biosystems qPCR Library Quantification Kit Illumina
  - [Accuprime Pfx Super Mix](http://tools.invitrogen.com/content/sfs/manuals/accuprimepfxsupermix_man.pdf)
  - [SequalPrep Normalization Plate (96) Kit](http://tools.lifetechnologies.com/content/sfs/manuals/sequalprep_platekit_man.pdf)
  - [Agilent High Sensitivity DNA Kit Guide](http://www.chem.agilent.com/library/usermanuals/Public/G2938-90321_SensitivityDNA_KG_EN.pdf)
  - [QIAquick Gel Extraction Kit](https://www.qiagen.com/us/resources/download.aspx?id=f4ba2d24-8218-452c-ad6f-1b6f43194425&lang=en)

***5.2) Initial Set up***

1. Reconstitute indexed primers and sequencing primers to 100 uM. See Appendix D for primer design.
2. Prepare 100ul 10 uM aliquots of indexed primers. Do not dilute sequencing primers.
3. Array aliquots of indexed primers into four 96 well plates using the following scheme:

a. A701 – A712 with A501 – A508 b. A701 – A712 with B501 – B508 c. B701 – B712 with B501 – B508 d. B701 – B712 with A501 – A508

Note: These primer plates can be stored at -20°C and used for subsequent runs.

1. Extract template DNA and array in 96 well format leaving two wells open. (One for a negative water control and another for the positive Mock Community control)
2. Using Illumina Experiment Manager, create a sample plate for each 96 well plate of template. Choose indexes that correspond to one of the four index pair plates above. See Appendix A for instruction on creating a custom assay in IEM.
3. Using Illumina Experiment Manager, create a sample sheet for the run. Ensure that index choices are compatible with one another and there is sufficient diversity in the index reads so as to activate both light channels every cycle. Note: Primer plate A has insufficient diversity to be run alone. If sequencing 96 or fewer samples, choose plate B, C, or D.

***5.3) PCR***

Note: These steps may be performed using an epMotion or similar automated pipetting system.

1. Dispense 17 ul of Accuprime Pfx Supermix into each well of a new 96 well plate.
2. Using a multichannel pipette, transfer 1 ul of template DNA per well to the corresponding well on the PCR plate.
3. Using a multichannel pipette, transfer 2 ul of each paired set of index primers from the primer plate to the corresponding well on the PCR plate. Be sure to follow the layout chosen in the sample sheet.
4. Add 1 ul of PCR grade H~2~O to the negative control well, and 1 ul of Mock Community at a 1:3 dilution to the positive control well.
5. Repeat for up to four 96 well plates. Seal plates, vortex briefly and spin down contents.
6. Place in thermocycler.

Use the following program:

95°C 2:00
--------30 cycles--------
95°C 00:20
55°C 00:15
72°C 5:00
----------------------
72°C 10:00
4°C Hold

***5.4) Gel Electrophoresis***

1. 1 or 2 random rows of 12 should be selected from each PCR plate and run on a gel to confirm success of the PCR. (Alternatively all samples can be run on a single E-Gel)
2. Use 2 ul of sample, 4 ul of loading dye in a 1% agarose gel.
3. Run at 100v for 30 minutes alongside a 1kb+ ladder.
4. Photograph gel under UV. Check to be sure there is a band for every well.

***5.6) Cleanup, Normalization, and Pooling***

Use the SequalPrep Normalization Plate Kit

1. Transfer 18 ul of PCR product from PCR plate to corresponding well on the normalization plate.
2. Add 18 ul of Binding Buffer. Mix by pipetting, sealing, vortexing, and spinning briefly.
3. Incubate at room temperature for 60 minutes. Plate can be incubated overnight at 4C if needed. Extra time does not improve results.
4. Aspirate the liquid from the wells. Do not scrape the sides.
5. Add 50 ul of Wash Buffer and pipette up and down twice, then aspirate immediately. Invert and tap plate on a paper towel to Ensure there is no residual wash buffer in any wells.
6. Add 20 ul of Elution Buffer. Mix by pipetting up and down 5 times. Seal, vortex, and spin briefly.
7. Incubate at room temperature for 5 minutes.
8. Create a pool from each plate. Take 5 ul of each well to pool. The use of an empty 96 well plate may facilitate the use of multichannel pipettes.
9. Freeze the remaining sample for later use.

***5.7) Library QC & Quantification***

1. Prepare the following dilutions of each pooled library in PCR grade H~2~O (or 10nM Tris-HCl + 0.05% Tween20):

a. 1:1

b. 1:10

c. 1:1000 (dilute in several steps for better results)

d. 1:2000

e. 1:4000

2. (Optional) Agilent Bioanalyzer Trace

a. Prepare Gel-Dye mix if not already prepared.

b. Let reagents equilibrate to room temperature.

c. Turn Bioanalyzer on and load 350 ul of dH~2~O onto electrode cleanser and place in analyzer for 5 minutes.

d. Open a high sensitivity chip and place on the priming station. Base plate should be a position “C” and syringe clip should be at lowest position.

e. Load 9.0 ul of gel-dye mix to position 12 market with a large “G”. Ensure the syringe plunger is at 1.0 ml and close the station. Press plunger until it is held by clip.

f. Wait for exactly 60 seconds then release the plunger clip. Wait an additional 5 seconds, then slowly pull the plunger back to the 1.0 ml position.

g. Open the priming station. Pipette 9.0 ul of gel-dye mix into the other wells marked “G” in positions 4,8,and 16.

h. Pipette 5.0 ul of marker to all wells excluding the right column. (No marker positions 4,8,12, and 16)

i. Load 1 ul of ladder into position 15 marked by the ladder symbol.

j. Pipette 1 ul of each of dilutions a – b above. Top row Plate 1 Pool 1:1 x 1, 1:10 x 2. Second row Plate 2 pool 1:1 x 1, 1:10 x 2. Third row Plate 3 Pool 1:1 x 1, 1:10 x 2. Bottom row Plate 4 Pool 1:1 x 1, 1:10 x 2.

k. Place chip in the designated vortex for 1 minute, then transfer chip to the Bioanalyzer.

l. Open the 2100 Expert software and select the HS DNA Assay. Enter sample names/dilutions for each of the test wells. Click Start.

m. Print .pdf when run finishes.

3. (Optional) Gel Purification

a. Load 50-200ul of each pooled plate on a 1% agarose gel. It will usually be necessary to tape several combs together to accomadate the volume.

b. Run the gel for ~1hr at 100V, until there is sufficient separation between the amplicon and the indices.

c. Briefly image the gel under UV light to locate and excise the band.

b. Follow the manufactures protocol for extracting the DNA from the gel ([QIAquick Gel Extraction Kit](https://www.qiagen.com/us/resources/download.aspx?id=f4ba2d24-8218-452c-ad6f-1b6f43194425&lang=en)).

c. Once the DNA is isolated, proceed to qPCR quantification.

4. Kapa qPCR Library Quantification

a. Before qPCR reaction setup, add 1 ml Primer Premix (10X) to the 5 ml bottle of KAPA SYBR® FAST qPCR Master Mix (2X) and mix by vortexing for 10 sec. Record the date of Primer Premix addition on the KAPA SYBR® FAST qPCR Master Mix bottle.

b. Reaction can be either 10 ul or 20 ul. A 10 ul reaction volume is recommended.

c. Prepare a 96 well qPCR plate compatible with the real time thermocycler. There are six standards. Each should be run in triplicate. Each pool at each dilution should be run in triplicate.

d. For 10 ul reaction volume dispense 6 ul of master mix into each well needed.

e. Pipette 4 ul of standards and library dilutions into appropriate wells. Mix by pipetting. Vortex and spin optional.

f. Place plate in thermocycler. Start control software

g. Program the following cycle

```
i.  Initial Activation 95°C 5 minutes

ii. 35 cycles

    1.  Denaturation 95°C 30 seconds

    2.  Annealing 60°C 45 seconds

    3.  If library fragment size exceeds 700bp, extend annealing
        step to 90 seconds.

iii. Perform melt curve to check for primer/adaptor dimer
```

h. Assign wells and group replicates.

i. Enter values for standards

```
i.  Std. 1 20pM

ii. Std. 2 2pM

iii. Std. 3 0.2pM

iv. Std. 4 0.02pM

v.  Std. 5 0.002pM

vi. Std. 6 0.0002pM

vii. Note: The concentrations provided here are for the DNA
    Standards as supplied in the kit, and are NOT the
    concentrations in the reactions. Provided that the volume of
    template added to each reaction is the same for Standards
    and for library samples (i.e. 4 ul in each case), there is
    no need to account for these volumes when calculating the
    concentrations of library samples, nor should one need to
    calculate the concentration of template in the reaction.
```

j. Run program

k. To calculate library concentration use the following formula:

```
i.  Average x (452/Avg fragment length from bioanalyzer) x
    dilution factor

ii. Use the average of the triplicate data points corresponding
    to the most concentrated library DNA dilution that falls
    within the dynamic range of the DNA Standards to calculate
    the concentration of the undiluted library.

iii. Do not include outliers in calculation. If there is more
    than one outlier in a group, the assay must be repeated.
```

5. Create a single final library by pooling each of the 4 pooled plates into a single well. Be sure to pool such that each plate has an equal final conentration (not necessarily equal volumes. Hint: C~1~ V~1~ = C~2~ V~2~ ). Final pool must be >10ul in total volume (40-80ul of >1nM library is ideal)

***5.8) Sequencing***

1. Remove a 500 cycle reagent cartridge from the -20°C freezer. Place in room temperature water bath for one hour. Place HT1 buffer tube in 4°C fridge. While reagent cartridge is thawing, perform steps 2-6.
2. Prepare fresh 0.2N NaOH.
3. To a 1.5ml tube, add 10 ul of library and 10 ul of 0.2N NaOH. To a separate tube add 2 ul PhiX, 3 ul PCR grade water, and 5 ul of 0.2N NaOH. Pipette to mix. Note: NaOH concentration on the flow cell must remain under 0.001N. Adjusting the concentration of the NaOH used to denature the DNA to 0.1N may be necessary if library concentration is 1nM or below.[^1]
4. Allow the tubes to incubate at room temperature for 5 minutes. Immediately add 980 ul of ice-cold HT1 to the library tube, and 990 ul HT1 to the PhiX tube. Note: the resulting 20pM PhiX can be frozen and used for subequent runs.
5. Use HT1 to further dilute both the library and PhiX to 4pM for a v2 kit. Can load up to 8pM for a v3 kit.

See example below: a. (1.45 nM library x 10 ul) + (0.2N NaOH x 10 ul) + 980 ul HT1 = 14.5pM Lib, 0.002N NaOH

b. (14.5pM lib x 275.86 ul) + 724.14 ul HT1 = 4.0pM lib, 0.00055N NaOH

c. [(10nM PhiX x 2 ul) + 3 ul H~2~O] + (0.2N NaOH x 5 ul) + 990 ul HT1 = 20pM PhiX, 0.001N NaOH

d. (20pM PhiX x 200 ul) + 800 ul HT1 = 4.0pM PhiX, 0.0002N NaOH

e. (4.0pM Lib x 900 ul) + (4.0pM PhiX x 100 ul) = solution loaded

f. Solution loaded is 4.0pM overall with a 3.6pM Library concentration, 0.4pM PhiX concentration, and 0.000515N NaOH

6. For a 10% PhiX run, combine 900 ul of 4.0pM Library and 100 ul PhiX in a final tube. Vortex.
7. When the reagent cartridge has thawed, dry bottom with paper towel. Invert the cartridge repeatedly to check each well is thawed. This also serves to mix the reagents. Place in hood.
8. Using a clean 1000 ul pipette tip, break the foil covering wells 12, 13, 14, and 17 of the reagent cartridge.
9. Load 600 ul of the final Libary/PhiX solution into well 17 on the reagent cartridge.
10. Place 3 ul of the 100 uM Read 1 Sequencing Primer(s) into a clean PCR tube. Repeat in separate tubes for the Index Primer(s) and Read 2 Sequencing Primer(s).
11. Use an extra long 100 ul tip and pipetter transfer the 3 ul of Read 1 Sequencing Primer to the bottom of well 12 and pipette to mix. Repeat this process spiking the Index Primer into well 13 and the Read 2 Sequencing Primer into well 14.
12. Set reagent cartridge aside. Unbox flow cell and PR2 bottle.
13. Thoroughly rinse the flow cell with Milli-Q water. Carefully dry by blotting with lint free wipes (Kimwipes). Give special attention to the edges and points of intersection between the glass and plastic.
14. Wet a new wipe with 100% alcohol and wipe the glass on both sides avoiding the rubber intake ports.
15. Visually inspect the flow cell to ensure there are no blemishes, particles, or fibers on the glass.
16. Transfer reagent cartridge, flow cell, PR2 bottle, and flash drive with the sample sheet to the MiSeq.
17. Copy Sample Sheet from the flash drive to the "Sample Sheets" folder on the desktop of the MiSeq
18. Follow on screen instructions to load the flow cell, reagent cartridge, and PR2 bottle. Empty and replace the waste bottle.
19. Ensure the machine recognizes the correct sample sheet and the run parameters are correct.
20. Wait for the MiSeq to perform its pre-run checks, and press start.

***5.9) Run Monitoring***

1. The run should be monitored periodically using Illumina Sequence Analysis Viewer.
2. Ideal parameters for a 90% 16S run:

a. Cluster density 700-800k/mm2 for v2 kits

b. Cluster density 1000-1100k/mm2 for v3 kits

c. >85% clusters passing filter

d. 10% aligned (amount of PhiX)

e. No spikes in corrected intensity plot

f. All indices identified following index reads

g. Final >Q30 score of >70%

***5.10) Final Steps***

1. Perform a post run wash on the MiSeq.
2. Dispose of liquid waste in appropriate hazardous jug and reagent cartridge in hazardous bucket.
3. When MiSeq Reporter finishes, copy the fastq files from the output folder to the run folder on the NAS drive.
4. Perform maintenance or standby wash if required.
5. Check data to confirm they are of sufficient quality and quantity.

**Appendix A: Adding An Assay To Illumina Experiment Manager**

Note: You can skip steps 2-9 by simply saving the [Schloss.txt](https://github.com/SchlossLab/MiSeq_WetLab_SOP/blob/master/Schloss.txt) file to the "C:\program files\Illumina\Illumina Experiment Manager\Sample Prep Kits" directory.

**Introduction**

Illumina Experiment Manager (IEM) is used to generate sample plates and sheets. A new assay must be added to the system to efficiently prepare sample sheets for 16S sequencing. Not only does this eliminate the need to manually assemble a sample sheet for 16S runs, but allows the user to retain use of the IEM index analysis feature. This ensures the indices selected for a particular run have sufficient diversity on a cycle-by-cycle basis and will successfully demultiplex.

Procedure

1. Open C:\program files\Illumina\Illumina Experiment Manager\Sample Prep Kits
2. Copy the file Nextera.txt, and rename the file Schloss.txt
3. Open in a text editor
4. Under [Name] change to Schloss
5. Under [PlateExtension] change to Schloss
6. Under [I7], clear the Nextera indices and paste in the SA701 through SB712 index names, and the REVERSE COMPLIMENT of the primer index sequences.
7. Under [I5], remove the Nextera indices and paste in the SA501 through SB512 index names and index sequences with no alteration (NOT the reverse compliment).
8. Under [DefaultLayout_SingleIndex] and [DefaultLayout_DualIndex] each well of a 96 well plate is listed with a corresponding Nextera index name. These must be replaced with Schloss index names. It is recommended that a text editor with column select capability be used to leave the index name number unchanged (i.e. 701, 702, etc.), while replacing the character ”N” for Nextera with “SA” for Schloss A.
9. Save and close.
10. Return to the IEM folder and open the Applications folder. Open and edit the file GenerateFASTQ.txt.
11. Add a line at to the bottom with the text “Schloss”

**Appendix B: Creating Sample Plates and a Sample Sheet in IEM**

***Part1: Creating Sample Plates***

1. Open IEM and click Create Sample Plate.
2. Select "Schloss" from the Sample Perp Kit Selection menu. Click Next.
3. Enter the plate name (e.g. projectName_plate1). Click Next.
4. Click the Plate tab for a 96-well plate format.
5. Copy and paste sample names from an excel file, csv, etc., or enter the names manually.
6. Select the appropriate I7 and I5 indices in the upper and left pull-down menus using the following scheme.
   a. Plate 1: A701 – A712 with A501 – A508 b. Plate 2: A701 – A712 with B501 – B508 c. Plate 3: B701 – B712 with B501 – B508 d. Plate 4: B701 – B712 with A501 – A508
   Note: You can select the first two indices, click and drag to highlight the remaing index spots, then click "fill right" or "fill down".
7. Click Finish, then Save.
8. Repeat steps 1-7 for all sample plates.

***Part2: Creating a Sample Sheet***

1. From the IEM main menu, select Create Sample Sheet.
2. On the Instrument Selction page, select the MiSeq and click Next.
3. On the MiSeq Application Selection page, click Other, Fastq Only, Next.
4. Enter the barcode for the MiSeq Reagent Kit being used for the run.
5. Select "Schloss" as the Sample Prep Kit.
6. Enter the Experiment Name, Investigator Name, and Description.
7. Change the number of cycles to 251 for both read 1 and 2. Click Next.
   Note: You should not need to check or uncheck any boxes on the right. They should remain unchanged after selecting the Schloss Sample Prep Kit.
8. On the Sample Selection page, uncheck the Maximize box in the upper right corner.
9. Click Select Plate in upper left, and select the appropriate sample plate file for plate 1.
10. Click Select All at the bottom, then Add Selected Samples.
11. Check the Sample Sheet Status on the right.
    Note: For plate 1, IEM may give a warning of low diversity. This will go away when more plates are added.
    As long as the status is not "Invalid", you may proceed.
12. Repeat steps 9-11 for the remaining sample plates.
13. Click Finish, then Save.
    Note: In most cases the Sample Sheet should be saved to a flash drive to be transfered to the MiSeq.

**Appendix C: Primer design**

Overall design considerations

- The sequencing primers must have a melting temperature near 65°C. This can be achieved by altering the pad sequence
- The index sequences must balance the number of bases at each position. The index sequences listed here have a 25% ATGC composition at each site. If you are going to cherry pick indices from the list, make sure that you have even representation.

Generic PCR primer design:

AATGATACGGCGACCACCGAGATCTACAC <i5><pad><link><16Sf> VX.N5??

CAAGCAGAAGACGGCATACGAGAT <i7><pad><link><16Sr> VX.N7??

Generic read 1 primer design

<pad><link><16Sf> VX.read1

Generic read 2 primer design

<pad><link><16Sr> VX.read2

Generic index read primer design

Reverse complement of (<pad><link><16Sr>) VX.p7_index

The listed sequences in the generic design, above, are the adapter sequences to allow annealing of the amplicons to the flow cell. The i5 and i7 sequences are the 8-nt index sequences. The pad is a 10-nt sequence to boost the sequencing primer melting temperatures. The link is a 2-nt sequence that is anti-complementary to the known sequences. The 16Sf and 16Sr are the gene specific primer sequences. Primers are purchased from IDT with no special purification. This system should work for any other region of the 16S rRNA gene or any other gene. The only thing to change would be the 16Sf/16Sr sequences and confirm that when combined with the pad sequence that the melting temperature is near 65°C.

16Sf

V3: CCTACGGGAGGCAGCAG

V4: GTGCCAGCMGCCGCGGTAA

16Sr

V4: GGACTACHVGGGTWTCTAAT

V5: CCCGTCAATTCMTTTRAGT

Link:

V4f: GT

V4r: CC

V3f: GG

V5r: GG

Pad:

Forward: TATGGTAATT

Reverse: AGTCAGTCAG

i5

SA501 ATCGTACG

SA502 ACTATCTG

SA503 TAGCGAGT

SA504 CTGCGTGT

SA505 TCATCGAG

SA506 CGTGAGTG

SA507 GGATATCT

SA508 GACACCGT

SB501 CTACTATA

SB502 CGTTACTA

SB503 AGAGTCAC

SB504 TACGAGAC

SB505 ACGTCTCG

SB506 TCGACGAG

SB507 GATCGTGT

SB508 GTCAGATA

SC501 ACGACGTG

SC502 ATATACAC

SC503 CGTCGCTA

SC504 CTAGAGCT

SC505 GCTCTAGT

SC506 GACACTGA

SC507 TGCGTACG

SC508 TAGTGTAG

SD501 AAGCAGCA

SD502 ACGCGTGA

SD503 CGATCTAC

SD504 TGCGTCAC

SD505 GTCTAGTG

SD506 CTAGTATG

SD507 GATAGCGT

SD508 TCTACACT

i7

SA701 AACTCTCG

SA702 ACTATGTC

SA703 AGTAGCGT

SA704 CAGTGAGT

SA705 CGTACTCA

SA706 CTACGCAG

SA707 GGAGACTA

SA708 GTCGCTCG

SA709 GTCGTAGT

SA710 TAGCAGAC

SA711 TCATAGAC

SA712 TCGCTATA

SB701 AAGTCGAG

SB702 ATACTTCG

SB703 AGCTGCTA

SB704 CATAGAGA

SB705 CGTAGATC

SB706 CTCGTTAC

SB707 GCGCACGT

SB708 GGTACTAT

SB709 GTATACGC

SB710 TACGAGCA

SB711 TCAGCGTT

SB712 TCGCTACG

SC701 ACCTACTG

SC702 AGCGCTAT

SC703 AGTCTAGA

SC704 CATGAGGA

SC705 CTAGCTCG

SC706 CTCTAGAG

SC707 GAGCTCAT

SC708 GGTATGCT

SC709 GTATGACG

SC710 TAGACTGA

SC711 TCACGATG

SC712 TCGAGCTC

SD701 ACCTAGTA

SD702 ACGTACGT

SD703 ATATCGCG

SD704 CACGATAG

SD705 CGTATCGC

SD706 CTGCGACT

SD707 GCTGTAAC

SD708 GGACGTTA

SD709 GGTCGTAG

SD710 TAAGTCTC

SD711 TACACAGT

SD712 TTGACGCA

Primers used to amplify 1536 samples using the V4 region. If you only want 384 then use a subset of the listed primers (e.g. all of the v4.SA5* and v4.SB5* and v4.SA7* and v4.SB7* primers):

v4.SA501 AATGATACGGCGACCACCGAGATCTACACATCGTACGTATGGTAATTGTGTGCCAGCMGCCGCGGTAA

v4.SA502 AATGATACGGCGACCACCGAGATCTACACACTATCTGTATGGTAATTGTGTGCCAGCMGCCGCGGTAA

v4.SA503 AATGATACGGCGACCACCGAGATCTACACTAGCGAGTTATGGTAATTGTGTGCCAGCMGCCGCGGTAA

v4.SA504 AATGATACGGCGACCACCGAGATCTACACCTGCGTGTTATGGTAATTGTGTGCCAGCMGCCGCGGTAA

v4.SA505 AATGATACGGCGACCACCGAGATCTACACTCATCGAGTATGGTAATTGTGTGCCAGCMGCCGCGGTAA

v4.SA506 AATGATACGGCGACCACCGAGATCTACACCGTGAGTGTATGGTAATTGTGTGCCAGCMGCCGCGGTAA

v4.SA507 AATGATACGGCGACCACCGAGATCTACACGGATATCTTATGGTAATTGTGTGCCAGCMGCCGCGGTAA

v4.SA508 AATGATACGGCGACCACCGAGATCTACACGACACCGTTATGGTAATTGTGTGCCAGCMGCCGCGGTAA

v4.SB501 AATGATACGGCGACCACCGAGATCTACACCTACTATATATGGTAATTGTGTGCCAGCMGCCGCGGTAA

v4.SB502 AATGATACGGCGACCACCGAGATCTACACCGTTACTATATGGTAATTGTGTGCCAGCMGCCGCGGTAA

v4.SB503 AATGATACGGCGACCACCGAGATCTACACAGAGTCACTATGGTAATTGTGTGCCAGCMGCCGCGGTAA

v4.SB504 AATGATACGGCGACCACCGAGATCTACACTACGAGACTATGGTAATTGTGTGCCAGCMGCCGCGGTAA

v4.SB505 AATGATACGGCGACCACCGAGATCTACACACGTCTCGTATGGTAATTGTGTGCCAGCMGCCGCGGTAA

v4.SB506 AATGATACGGCGACCACCGAGATCTACACTCGACGAGTATGGTAATTGTGTGCCAGCMGCCGCGGTAA

v4.SB507 AATGATACGGCGACCACCGAGATCTACACGATCGTGTTATGGTAATTGTGTGCCAGCMGCCGCGGTAA

v4.SB508 AATGATACGGCGACCACCGAGATCTACACGTCAGATATATGGTAATTGTGTGCCAGCMGCCGCGGTAA

v4.SC501 AATGATACGGCGACCACCGAGATCTACACACGACGTGTATGGTAATTGTGTGCCAGCMGCCGCGGTAA v4.SC502 AATGATACGGCGACCACCGAGATCTACACATATACACTATGGTAATTGTGTGCCAGCMGCCGCGGTAA v4.SC503 AATGATACGGCGACCACCGAGATCTACACCGTCGCTATATGGTAATTGTGTGCCAGCMGCCGCGGTAA v4.SC504 AATGATACGGCGACCACCGAGATCTACACCTAGAGCTTATGGTAATTGTGTGCCAGCMGCCGCGGTAA v4.SC505 AATGATACGGCGACCACCGAGATCTACACGCTCTAGTTATGGTAATTGTGTGCCAGCMGCCGCGGTAA v4.SC506 AATGATACGGCGACCACCGAGATCTACACGACACTGATATGGTAATTGTGTGCCAGCMGCCGCGGTAA v4.SC507 AATGATACGGCGACCACCGAGATCTACACTGCGTACGTATGGTAATTGTGTGCCAGCMGCCGCGGTAA v4.SC508 AATGATACGGCGACCACCGAGATCTACACTAGTGTAGTATGGTAATTGTGTGCCAGCMGCCGCGGTAA v4.SD501 AATGATACGGCGACCACCGAGATCTACACAAGCAGCATATGGTAATTGTGTGCCAGCMGCCGCGGTAA v4.SD502 AATGATACGGCGACCACCGAGATCTACACACGCGTGATATGGTAATTGTGTGCCAGCMGCCGCGGTAA v4.SD503 AATGATACGGCGACCACCGAGATCTACACCGATCTACTATGGTAATTGTGTGCCAGCMGCCGCGGTAA v4.SD504 AATGATACGGCGACCACCGAGATCTACACTGCGTCACTATGGTAATTGTGTGCCAGCMGCCGCGGTAA v4.SD505 AATGATACGGCGACCACCGAGATCTACACGTCTAGTGTATGGTAATTGTGTGCCAGCMGCCGCGGTAA v4.SD506 AATGATACGGCGACCACCGAGATCTACACCTAGTATGTATGGTAATTGTGTGCCAGCMGCCGCGGTAA v4.SD507 AATGATACGGCGACCACCGAGATCTACACGATAGCGTTATGGTAATTGTGTGCCAGCMGCCGCGGTAA v4.SD508 AATGATACGGCGACCACCGAGATCTACACTCTACACTTATGGTAATTGTGTGCCAGCMGCCGCGGTAA

v4.SA701 CAAGCAGAAGACGGCATACGAGATAACTCTCGAGTCAGTCAGCCGGACTACHVGGGTWTCTAAT

v4.SA702 CAAGCAGAAGACGGCATACGAGATACTATGTCAGTCAGTCAGCCGGACTACHVGGGTWTCTAAT

v4.SA703 CAAGCAGAAGACGGCATACGAGATAGTAGCGTAGTCAGTCAGCCGGACTACHVGGGTWTCTAAT

v4.SA704 CAAGCAGAAGACGGCATACGAGATCAGTGAGTAGTCAGTCAGCCGGACTACHVGGGTWTCTAAT

v4.SA705 CAAGCAGAAGACGGCATACGAGATCGTACTCAAGTCAGTCAGCCGGACTACHVGGGTWTCTAAT

v4.SA706 CAAGCAGAAGACGGCATACGAGATCTACGCAGAGTCAGTCAGCCGGACTACHVGGGTWTCTAAT

v4.SA707 CAAGCAGAAGACGGCATACGAGATGGAGACTAAGTCAGTCAGCCGGACTACHVGGGTWTCTAAT

v4.SA708 CAAGCAGAAGACGGCATACGAGATGTCGCTCGAGTCAGTCAGCCGGACTACHVGGGTWTCTAAT

v4.SA709 CAAGCAGAAGACGGCATACGAGATGTCGTAGTAGTCAGTCAGCCGGACTACHVGGGTWTCTAAT

v4.SA710 CAAGCAGAAGACGGCATACGAGATTAGCAGACAGTCAGTCAGCCGGACTACHVGGGTWTCTAAT

v4.SA711 CAAGCAGAAGACGGCATACGAGATTCATAGACAGTCAGTCAGCCGGACTACHVGGGTWTCTAAT

v4.SA712 CAAGCAGAAGACGGCATACGAGATTCGCTATAAGTCAGTCAGCCGGACTACHVGGGTWTCTAAT

v4.SB701 CAAGCAGAAGACGGCATACGAGATAAGTCGAGAGTCAGTCAGCCGGACTACHVGGGTWTCTAAT

v4.SB702 CAAGCAGAAGACGGCATACGAGATATACTTCGAGTCAGTCAGCCGGACTACHVGGGTWTCTAAT

v4.SB703 CAAGCAGAAGACGGCATACGAGATAGCTGCTAAGTCAGTCAGCCGGACTACHVGGGTWTCTAAT

v4.SB704 CAAGCAGAAGACGGCATACGAGATCATAGAGAAGTCAGTCAGCCGGACTACHVGGGTWTCTAAT

v4.SB705 CAAGCAGAAGACGGCATACGAGATCGTAGATCAGTCAGTCAGCCGGACTACHVGGGTWTCTAAT

v4.SB706 CAAGCAGAAGACGGCATACGAGATCTCGTTACAGTCAGTCAGCCGGACTACHVGGGTWTCTAAT

v4.SB707 CAAGCAGAAGACGGCATACGAGATGCGCACGTAGTCAGTCAGCCGGACTACHVGGGTWTCTAAT

v4.SB708 CAAGCAGAAGACGGCATACGAGATGGTACTATAGTCAGTCAGCCGGACTACHVGGGTWTCTAAT

v4.SB709 CAAGCAGAAGACGGCATACGAGATGTATACGCAGTCAGTCAGCCGGACTACHVGGGTWTCTAAT

v4.SB710 CAAGCAGAAGACGGCATACGAGATTACGAGCAAGTCAGTCAGCCGGACTACHVGGGTWTCTAAT

v4.SB711 CAAGCAGAAGACGGCATACGAGATTCAGCGTTAGTCAGTCAGCCGGACTACHVGGGTWTCTAAT

v4.SB712 CAAGCAGAAGACGGCATACGAGATTCGCTACGAGTCAGTCAGCCGGACTACHVGGGTWTCTAAT

v4.SC701 CAAGCAGAAGACGGCATACGAGATACCTACTGAGTCAGTCAGCCGGACTACHVGGGTWTCTAAT v4.SC702 CAAGCAGAAGACGGCATACGAGATAGCGCTATAGTCAGTCAGCCGGACTACHVGGGTWTCTAAT v4.SC703 CAAGCAGAAGACGGCATACGAGATAGTCTAGAAGTCAGTCAGCCGGACTACHVGGGTWTCTAAT v4.SC704 CAAGCAGAAGACGGCATACGAGATCATGAGGAAGTCAGTCAGCCGGACTACHVGGGTWTCTAAT v4.SC705 CAAGCAGAAGACGGCATACGAGATCTAGCTCGAGTCAGTCAGCCGGACTACHVGGGTWTCTAAT v4.SC706 CAAGCAGAAGACGGCATACGAGATCTCTAGAGAGTCAGTCAGCCGGACTACHVGGGTWTCTAAT v4.SC707 CAAGCAGAAGACGGCATACGAGATGAGCTCATAGTCAGTCAGCCGGACTACHVGGGTWTCTAAT v4.SC708 CAAGCAGAAGACGGCATACGAGATGGTATGCTAGTCAGTCAGCCGGACTACHVGGGTWTCTAAT v4.SC709 CAAGCAGAAGACGGCATACGAGATGTATGACGAGTCAGTCAGCCGGACTACHVGGGTWTCTAAT v4.SC710 CAAGCAGAAGACGGCATACGAGATTAGACTGAAGTCAGTCAGCCGGACTACHVGGGTWTCTAAT v4.SC711 CAAGCAGAAGACGGCATACGAGATTCACGATGAGTCAGTCAGCCGGACTACHVGGGTWTCTAAT v4.SC712 CAAGCAGAAGACGGCATACGAGATTCGAGCTCAGTCAGTCAGCCGGACTACHVGGGTWTCTAAT v4.SD701 CAAGCAGAAGACGGCATACGAGATACCTAGTAAGTCAGTCAGCCGGACTACHVGGGTWTCTAAT v4.SD702 CAAGCAGAAGACGGCATACGAGATACGTACGTAGTCAGTCAGCCGGACTACHVGGGTWTCTAAT v4.SD703 CAAGCAGAAGACGGCATACGAGATATATCGCGAGTCAGTCAGCCGGACTACHVGGGTWTCTAAT v4.SD704 CAAGCAGAAGACGGCATACGAGATCACGATAGAGTCAGTCAGCCGGACTACHVGGGTWTCTAAT v4.SD705 CAAGCAGAAGACGGCATACGAGATCGTATCGCAGTCAGTCAGCCGGACTACHVGGGTWTCTAAT v4.SD706 CAAGCAGAAGACGGCATACGAGATCTGCGACTAGTCAGTCAGCCGGACTACHVGGGTWTCTAAT v4.SD707 CAAGCAGAAGACGGCATACGAGATGCTGTAACAGTCAGTCAGCCGGACTACHVGGGTWTCTAAT v4.SD708 CAAGCAGAAGACGGCATACGAGATGGACGTTAAGTCAGTCAGCCGGACTACHVGGGTWTCTAAT v4.SD709 CAAGCAGAAGACGGCATACGAGATGGTCGTAGAGTCAGTCAGCCGGACTACHVGGGTWTCTAAT v4.SD710 CAAGCAGAAGACGGCATACGAGATTAAGTCTCAGTCAGTCAGCCGGACTACHVGGGTWTCTAAT v4.SD711 CAAGCAGAAGACGGCATACGAGATTACACAGTAGTCAGTCAGCCGGACTACHVGGGTWTCTAAT v4.SD712 CAAGCAGAAGACGGCATACGAGATTTGACGCAAGTCAGTCAGCCGGACTACHVGGGTWTCTAAT

Read 1 primer for V4 region

TATGGTAATTGTGTGCCAGCMGCCGCGGTAA

Read 2 primer for V4 region

AGTCAGTCAGCCGGACTACHVGGGTWTCTAAT

Index primer for V4 region

ATTAGAWACCCBDGTAGTCCGGCTGACTGACT

[^1]: For extremely low concentration libraries we have adapted a method published by Quail et al.[http://www.nature.com/nmeth/journal/v5/n12/full/nmeth.1270.html](https://github.com/SchlossLab/MiSeq_WetLab_SOP/blob/customXml/item1.xml)

Libraries as low as 0.1nM do not allow for sufficient dilution to

reduce NaOH to 0.001N. It must be neutralized using 200mM Tris pH

7.0. Example: 40 ul 0.1nM library + 40 ul 0.1N NaOH. Incubate for 5

min. Add 80 ul 200mM Tris. Then add 840 ul HT1. This results in a

4.0pM library.
